# Autologous micrograft accelerates endogenous wound healing response through ERK-induced cell migration

**DOI:** 10.1101/545376

**Authors:** Martina Balli, Francesca Vitali, Adrian Janiszewski, Ellen Caluwé, Alvaro Cortés-Calabuig, Robin Duelen, Flavio Ronzoni, Riccardo Bellazzi, Aernout Luttun, Maria G. Cusella De Angelis, Gabriele Ceccarelli, Frederic Lluis, Maurilio Sampaolesi

## Abstract

Defective fibroblast migration causes delayed wound healing (WH) and chronic skin lesions. Autologous micrograft (AMG) therapies have recently emerged as a new effective treatment able to improve wound healing capacity. However, the molecular mechanisms connecting their beneficial outcomes with the wound healing process are still unrevealed. Here, we show that AMG modulates primary fibroblast migration and accelerates skin re-epithelialization without affecting cell proliferation. We demonstrate that AMG is enriched in a pool of WH-associated growth factors that may provide the initiation signal for a faster endogenous wound healing response. This, in turn leads to increased cell migration rate by elevating activity of extracellular signal-regulated kinase (ERK) pathway and subsequent activation of matrix metalloproteinase expression and their extracellular enzymatic activity. Moreover, AMG-treated wounds showed increased granulation tissue formation and organized collagen content. Overall, we shed light on AMG molecular mechanism supporting its potential to trigger a highly improved wound healing process.

## Introduction

Traumatic skin injuries and impaired wound healing conditions lead to a widespread public health and economical problem, hindering patient quality of life, who require long-term hospitalization, finally contributing to mortality.

Autologous micrograft (AMG) technique is an emerging treatment for skin damages that has demonstrated to improve WH, minimize scar formation and is an affordable alternative to traditional skin grafting. AMG treatment has been reported to ameliorate the repair of dehiscent wounds (*1*) and traumatic wounds associated with hypertrophic skin/cardiac scars (*2, 3*). An important advantage of AMG treatment is the use of micrograft drawn directly from the subject, thus limiting rejection and allowing wound coverage by using a minimal amount of donor tissue. However, currently, whether AMG performs uniquely a passive protective function by wound coverage or in addition actively promotes wound healing response is not completely understood. Furthermore, the putative molecular mechanisms activated upon AMG treatment on receptor-damaged cells have previously never been studied.

Wound healing is a well-orchestrated process that involves the sequential collaborative effort of different players such as several cell types, extracellular matrix (ECM) components, growth factors and signaling pathways. In the first stage of WH, cytokines and growth factors regulate the inflammatory response leading to the initiation of the wound repair process. Several studies have provided the list of fundamental growth factors that trigger initial WH response such as epidermal growth factor (EGF), basic fibroblast growth factor (bFGF), insulin-like growth factor (IGF), platelet-derived growth factor (PDGF) and transforming growth factor beta (TGF-β) (*4*). Next, growth factor activation of intracellular signaling pathways, such as the Mitogen Activated Protein Kinase (MAPK) signaling, promotes the progression to the next essential WH stages such as fibroblast proliferation, migration and differentiation into myofibroblasts (*5*). Fibroblast are also responsible for the secretion of extracellular enzymes, such as the matrix metalloproteinases (MMPs) (*6*), that will allow ECM remodeling by cell migration, re-epithelialization and finally facilitating wound closure.

The MMPs belong to a zinc-containing endopeptidase family which consists of 25 proteases in humans and 24 in mice (*6*) that directly affect cell–matrix adhesion and ECM remodeling (*7*). MMP-1a, MMP3 and MMP9 are known as major chemokine regulators during WH (*7*), while fibroblast expression of Mmp13 regulates myofibroblast functions and granulation tissue formation (*8*). Interestingly, *Mmp3*- and *Mmp13*-deficient mice showed retarded WH and wound contraction failure, demonstrating the essential role of MMP family members during the WH response (*9, 10*). The transcriptional regulation of MMPs is partially executed by the activity of activator protein-1 (AP-1) transcription factor (*11*). Increased JUN and FOS protein levels trigger up-regulation of AP-1 target genes, such as MMPs. Interestingly, EGF, bFGF, TGF-β and tumour necrosis factor-α (TNF-α) growth factors are able to stimulate AP-1 activity by phosphorylation of JUN or FOS proteins through MAPK signaling pathway activation (*11*). In agreement, *junb* deficiency leads to delayed tissue remodeling during WH (*12*).

Modulation of the extracellular signal-regulated kinase (ERK) - signaling pathway, a MAPK subgroup, leads to changes in cell proliferation, differentiation, inflammation and cell migration (*13*). ERK signaling activity is needed during WH (*14*) however, its specific downstream function is still controversial: while in some studies the acceleration of cutaneous WH via MAPK cascade activation has been attributed to increased cell proliferation (*15*), other studies have shown a specific role in cell migration induction (*14*).

In this work, we shed light on the molecular mechanisms of AMG-based treatment. We provide evidence that AMG treatment enhances *in vivo* wound closure, re-epithelialization and angiogenesis. We show that AMG solution is enriched in a myriad of pro-motility growth factors implicated in WH response initiation including bFGF, EGF and IGF-1. Furthermore, by comparative transcriptome analysis followed by Protein-Protein Interaction (PPI) network analysis of AMG-treated primary fibroblasts, we identified cell migration and ERK/MAPK signaling pathway as essential factors in the AMG-mediated WH response. We demonstrate that AMG treatment enhances ERK-dependent MMP gene expression and extracellular enzymatic activity in fibroblasts leading to *in vitro* scratch closure acceleration and speeds up WH process *in vivo* without affecting cell proliferation. Accordingly, inhibition of either ERK phosphorylation or MMP enzymatic activity significantly reverses the AMG-mediated WH effects. Therefore, our study reveals the molecular mechanisms of AMG-based treatment, and thus provides a groundwork for future advances in the therapeutic use of micrografts for WH.

## Results

### Wound-induced transcriptional network upon AMG treatment

Fibroblasts are determinant in supporting WH due to their property to migrate from the dermis to the wound site in response to the cytokine/growth factor gradient. Fibroblasts are also responsible for the production of bio-molecules that will allow ECM remodeling and support migratory cell abilities during the re-epithelialization phase (*4*) and angiogenesis (*16*). Inefficient fibroblast migration and function is a cause of impaired wound healing (*17*) underlying the critical role of fibroblasts during WH.

On the way towards the understanding of the molecular mechanism of AMG-mediated WH, we first followed a broad transcriptome analysis of AMG-treated adult primary fibroblasts. Firstly, we designed a pilot study to explore how AMG treatment impacts the transcriptome of wounded fibroblasts over extended time and to select a potentially most relevant time point for further in-depth analysis of direct response to AMG treatment. For that we performed an *in vitro* scratch assay using primary mouse adult fibroblasts. Cells were subsequently treated with AMG for 1h, 5h, 12h and 24h and together with untreated wounded fibroblasts as a control collected for RNA-seq analysis (deposited in GEO Datasets under accession no. GSE123829). We took advantage of the N-of-1 Pathway MixEnrich (*18*) analysis, allowing the identification of differentially expressed genes (DEGs) without the requirement of large cohorts or replicates (Supplementary material and methods text) (fig. S1A). The pilot analysis shows that both AMG treatment and time of treatment have an effect on the transcriptome profile, as shown by the principal component analysis (fig. S1B). Interestingly, the 5h time point showed an enrichment of 40 gene ontology (GO) WH-related signaling pathways, such as positive regulation of cell motility, response to external stimulus, inflammatory response and ERK/MAPK cascade. In contrast, 12h and 24h AMG treatments showed an enrichment of 482 and 581 GO pathways, respectively, suggesting an accumulation of non-direct AMG-effects after 12h of treatment. We conclude that 5 hours of treatment likely best represents the earliest direct effects at the transcriptome level. We therefore focused further analyses on this time point (table S1; fig. S1C and fig. S1D).

To deeply evaluate the impact of AMG treatment at 5 hours, we expanded our biological cohort to n=3 by performing additional RNA-seq experiments. Differentially expressed genes (DEGs) between treated and untreated cells were then employed to conduct a double analysis strategy based on (i) conventional transcriptome analysis of DEG and (ii) Protein-Protein Interaction network-based analysis, both followed by enrichment analysis (Fig. 1A). Conventional transcriptome analytics showed 203 DEGs (Table S2) and 548 significantly enriched biological processes obtained using GO terms (GO-BP) (*19*) (GO enrichment analysis; Table S2) between AMG treated and untreated samples. GO-BP revealed involvement of important WH-associated biological processes and signaling pathways, such as cell migration, immune response, angiogenesis and MAPK cascade (fig. S1F and S1H; Table S2). Moreover, several genes, such as cytokines and MMPs, crucial for cell migration and WH were significantly differentially expressed (table S2; fig. S1G and S1I). This supports that AMG treatment rapidly induces changes in expression of genes associated with a successful wound healing process.

**Fig. 1.**
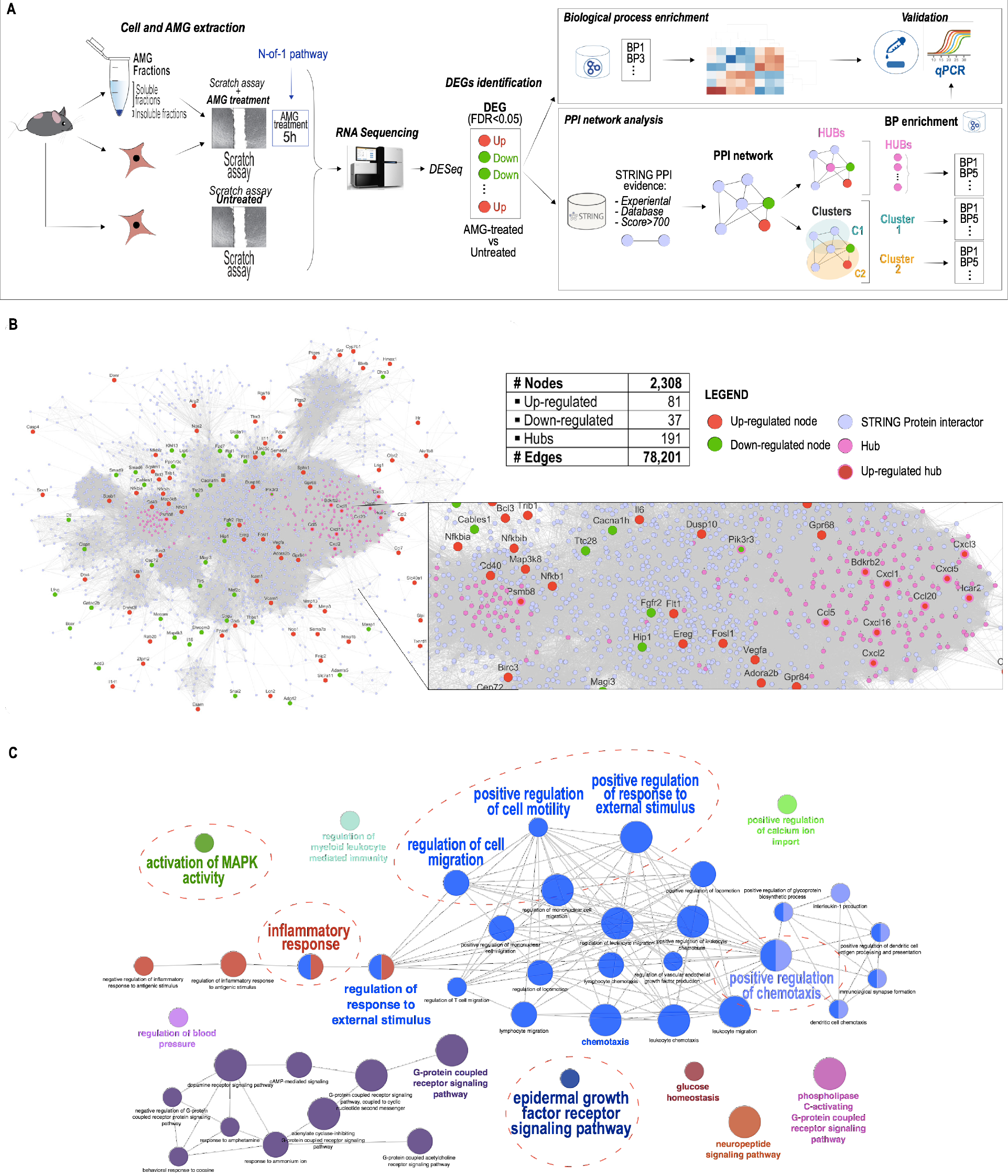
Comparative transcriptome analysis of primary murine fibroblasts upon AMG treatment. **(A)** Schematic representation of the strategy followed to reveal AMG-dependent mechanism on fibroblasts. Primary adult fibroblasts were obtained from murine tail’s tips and cultured until full confluency. A scratch wound assay was then performed using a pipette tip, creating an artificial wound in the cell monolayer. Autologous Micrograft (AMG) solutions were obtained starting from skin biopsies (1 cm^2^), through a method of disintegration using a sterile tissue destroyer. 1 mL of AMG solution was diluted 1:10 and applied. Transcriptome analysis via RNA-seq of AMG-treated and untreated cells led to the identification of AMG-related DEGs. Those genes were then used to perform a double strategy analysis: 1) Enrichment analysis of biological processes through GO and 2) PPI network analysis to predict DEGs-associated proteins and signaling pathways involved in the AMG molecular mechanism. **(B)** The Protein-Protein Interaction Network (PPI) referred to the list of differentially expressed genes (DEGs) observed upon AMG treatment and obtained by the STRING repository. DEGs are represented as red and green nodes of the network, based on their up-regulated or down-regulated expression, respectively. Hub nodes are presented as purple nodes, shown in an enlarged view. **(C)** Hub related GO enrichment analysis with highlighted biological process playing essential roles in the WH process.

For a deeper understanding of the mechanisms involved in AMG-wound healing process, we next constructed a PPI network starting from the identified DEGs and using the public PPI repository STRING (*20*) (Supplementary material – Network construction). The resulting network consisted of 2,308 protein nodes and 78,201 PPI edges (Fig. 1B; table S3). Through topological analysis of the network we derived 191 hub nodes (Supplementary material – Hub nodes; table S3), i.e. key nodes, and 49 significant network clusters (Supplementary material – clusters; table S3, fig. S2A), i.e. group of similar nodes grouped together. Interestingly, within the top 10 most significant clusters after enrichment analyses, we found 4 clusters (Cluster 1, 2, 6 and 7; fig. S2A) to be associated to WH and tissue regeneration processes such as ‘regulation of inflammatory response’ (fig. S2B), ‘regulation of cell migration/cell motility’ and ‘angiogenesis’ (fig. S2C), ‘activation of MAPK activity’ and ‘positive regulation of ERK1/ERK2 cascade’ (fig. S2D) as well as ‘wound healing’ (fig. S2E) in line with our conventional transcriptome analysis.

Both analysis strategies allowed us to conclude that AMG treatment might perform an active function on fibroblasts by stimulating several crucial WH-associated biological processes such as cell motility, cell migration and activation of MAPK signaling pathway.

### AMG treatment increases cell migration and accelerates *in vitro* scratch closure

To functionally evaluate the AMG effects on cellular functions, murine primary adult fibroblasts were isolated from tail’s tips and cultured to perform scratch wound assays. Scratched cells were then exposed to AMG treatment during different treatment times (1h, 5h, 12h and 24h). We found that, cells treated with AMG for 5 and 12 hours exhibited the quickest closure of the scratch wounds (Fig. 2A) with a closure percentage of 77% ± 11% and 84% ± 8% respectively at 24 hours compared to untreated cells (48% ± 26%). This shows that, independently of the treatment time, AMG increases cell migration rate and accelerates scratch closure compared to untreated condition.

**Fig. 2.**
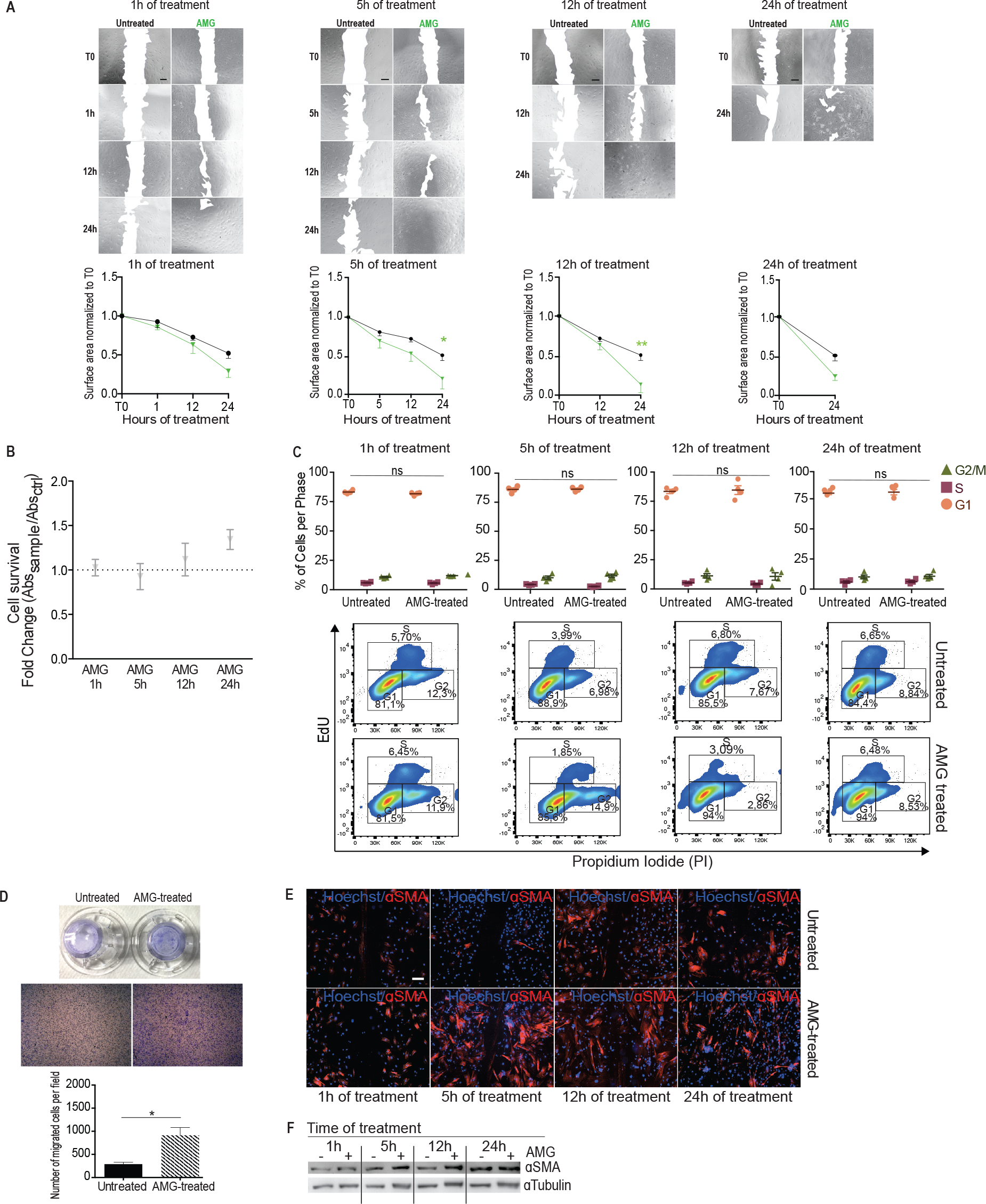
AMG treatment increases migration capacity of murine primary fibroblasts. **(A)** Representative images (up) and quantification (down) of AMG-treated and untreated fibroblasts in a scratch wound assay. AMG was applied for different time periods (1, 5, 12, 24h). Images were taken at the beginning (T0) and at regular intervals until closure was achieved. Quantification average of 4 biological replicates for each time of treatment are reported below. Data are presented as mean ± SEM (standard error of the mean). Significant differences vs control are calculated with two-way ANOVA and indicated as * P < 0.05; ** P < 0.01. **(B)** XTT assays were used to measure cell viability upon AMG-based treatment. Absorbance values (read at 450 nm) were normalized to the control group (untreated) and expressed as a fold change (n=5 biological replicates). **(C)** Cell cycle FACS analyses of AMG-treated fibroblasts for 1h, 5h, 12h and 24h. EdU and PI stainings were performed to detect cell cycle changes. Quantification average of 4 biological replicates is reported. Data are presented as mean ± SEM. **(D)** Macro- and microscopic observation of untreated and AMG-treated transwell chambers. Migrated cells were stained with crystal violet (0.1%), observed under a light microscope and analyzed using ImageJ software. Data are presented as mean ± SEM. P-values were calculated using student t test (4 biological replicates) * P < 0.05. **(E)** Representative immunofluorescence for ɑ-SMA protein localization (in red). Nuclei were stained with Hoechst (blue). Scale bar = 200 μm. **(F)** Representative western blotting showing ɑ-SMA protein level extracted from AMG-treated (+) or untreated (−) wounded murine fibroblasts.

To determine whether the enhanced wound closure is due to increased cell proliferation, cell viability and/or cell migration, we examined the effects of AMG treatment on cell survival by XTT assay and on cell cycle changes through EdU - 5-ethynyl-2’-deoxyuridine - assay. Interestingly, AMG treatment did not affect cell viability (Fig. 2B). In addition, cell cycle analysis showed no significant differences in any of the exanimated cell cycle stages (G0/G1, S and G2/M) between treated or untreated groups (Fig. 2C). Overall, we dismiss any possible effect on cell survival or on cell proliferation during AMG-dependent accelerated scratch closure.

We then analyzed the ability of the AMG treatment to regulate directed cell migration by using the transwell migration assay. Remarkably, AMG-treated cells showed a significant increase in the migration rate upon AMG treatment compared to untreated cells (Fig. 2D). In accordance, AMG led to the enrichment of alpha-Smooth Muscle Actin (*ɑ-SMA*) protein, a marker of activated contractile fibroblasts (also known as myofibroblasts; Fig. 2E). Western blotting confirmed that *ɑ-SMA* expression is enhanced in AMG-treated cells (Fig. 2F).

In conclusion, these results indicate that AMG treatment does not involve cell proliferation or survival functions in the accelerated scratch closure, but, instead, increases cell motility and accelerates *in vitro* wound closure.

### The soluble AMG fraction accelerates scratch wound closure compared to unprocessed micrografts

To gain knowledge about the mechanism by which AMG treatment accelerates *in vitro* scratch wound closure and enhances cell migration, we focused our attention on the components present within the AMG solution. Previously, studies suggested a putative positive role of mesenchymal stem cells (MSCs)-present in the AMG solution on WH (*2, 21*). However, a comprehensive study of the effects induced by different AMG fractions has been never performed.

Using a polyethylene glycol (PEG)-based membrane enrichment method, we separated the soluble fraction containing essential growth factors and/or cytokines from the insoluble fraction containing intra and extra membrane cellular vesicles. Next, wounded adult fibroblasts were either treated with unprocessed AMG, enriched-insoluble AMG fraction or enriched-soluble AMG fractions for 5h in an *in vitro* scratch wound assay. At 5h after treatment, untreated or treated samples with unprocessed or insoluble AMG fraction showed approximately 50% and 60% of wound closure with non-significant differences between conditions. In contrast, soluble AMG fraction treatment showed the 84% of wound closure with complete closure 12h after *in vitro* wounding. This shows that soluble AMG fraction significantly accelerates *in vitro* scratch closure compared to unprocessed or insoluble AMG treatment (Fig. 3A and 3B). Soluble fraction also granted enhanced cell migration capacity compared to unprocessed AMG confirmed by transwell migration assay (Fig. 3C).

**Fig. 3.**
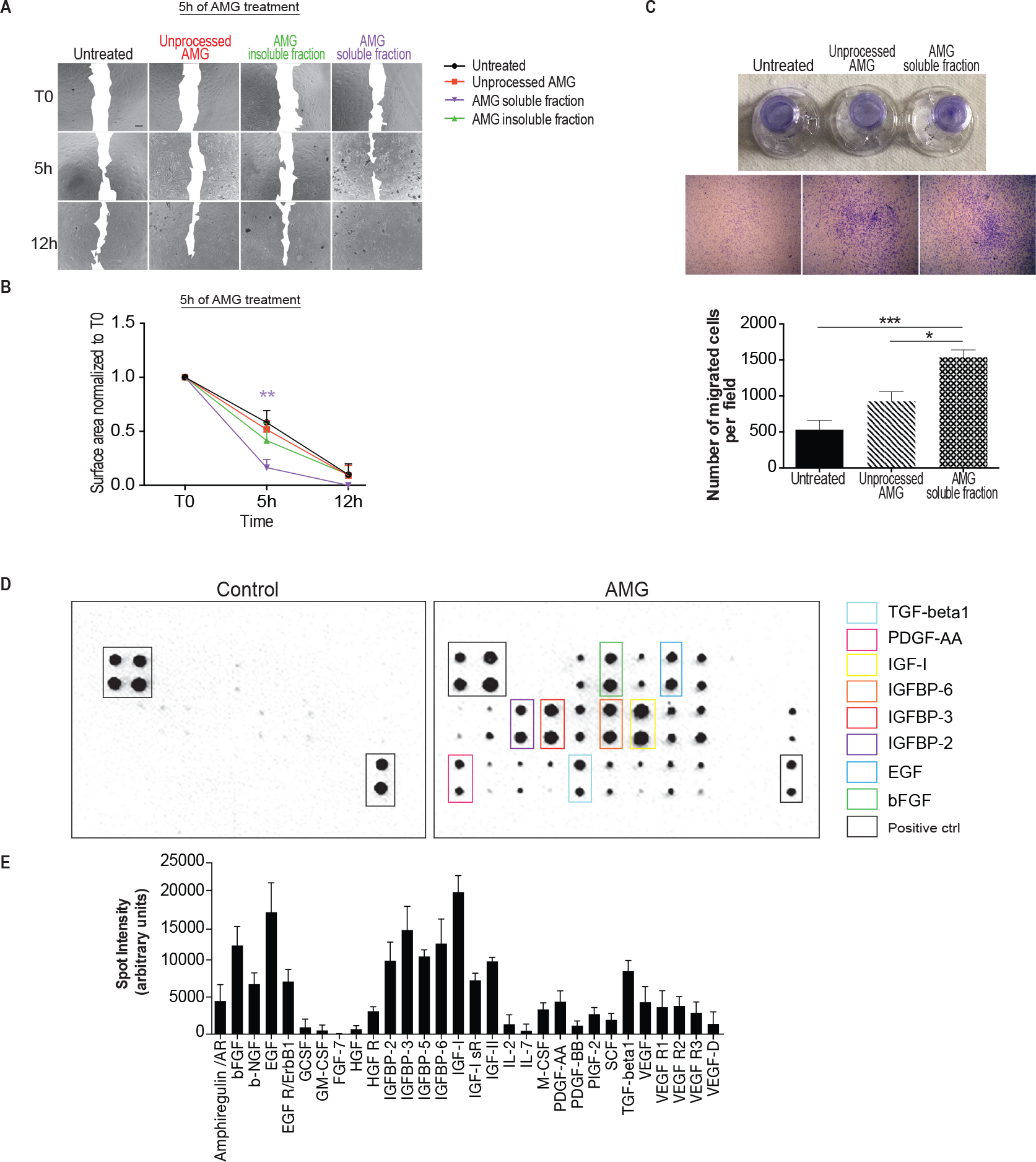
Characterization of AMG molecular fractions. **(A)** Representative image of scratch assay on AMG treated (5h) murine primary fibroblasts (unprocessed, insoluble and soluble AMG fractions). Scratched untreated cells were used as a control. Images were taken at the beginning (T0) and at regular intervals until closure was achieved. **(B)** Quantification average for each time of treatment. Data are presented as mean ± SEM. Differences are calculated with two-way ANOVA (3 biological replicates) and indicated as ** P < 0.01. **(C)** Observation of transwell chambers after stimulation with the AMG treatment. Migrated cells were stained with crystal violet 0.1%, observed under a light microscope and analyzed using ImageJ software. Data were presented as mean ± SEM. Differences are calculated using one-way ANOVA followed by Tukey test (4 biological replicates) and indicated as * P < 0.05; *** P < 0.005. **(D)** Representative images of mouse antibody array membrane showing presence of several growth factors within the AMG soluble fraction (right panel) compared to the PBS vehicle (left panel) (4 biological replicates). **(E)** Analysis of the increase in spot intensity (in arbitrary units) is shown in the graph.

To further characterize the factors contained in AMG solution, we analyzed the levels of growth factors within the AMG soluble fraction by using a growth factor antibody array. Interestingly, several members of IGF family, such as IGF-1, IGFBP-2, IGFBP-3 and IGFBP-6 as well as EGF, bFGF and TGF- β1 were highly detected within the AMG soluble fraction (Fig. 3D and 3E). Among those, EGF, bFGF and IGF have been implicated in triggering cell motility and migration (*4*). These findings indicate that AMG solutions are enriched in a myriad of pro-motility growth factors, which in turn may serve as inducers of the AMG-mediated increased cell migration capacity. Therefore, these finding may help to better understand the AMG composition and, especially, may provide a potential way to improve the regenerative effect of AMG.

### MMPs trigger AMG-mediated cell migration capacity

On the basis of the results above, we hypothesized that the growth factors present within the AMG solution are responsible for the activation of the pro-motility transcriptional program previously identified by PPI and GO enrichment analyses (Fig. 1C; fig. S1F; S1H; S2C, S2D). To further characterize key downstream factors involved in AMG-mediated acceleration of WH, we investigated the impact of AMG treatment on the transcriptome of wounded fibroblasts (Fig. 4A). We found that AMG induced significant changes in the expression 203 genes (log_2_FC >1, FDR<0.05). Among all differentially expressed genes, we identified several up-regulated members of the MMP family in the AMG treated conditions. We additionally validated these changes by qPCR. Among those, we validated the gene expression of *Mmp1a*, *Mmp1b* (known as collagenase-1), *Mmp9* (known as gelatinase B), *Mmp10* (stromelysin-2), *Mmp12* (metalloelastase) and *Mmp13*, confirming their significant up-regulated expression upon AMG treatment (Fig. 4B). MMPs are key regulators of ECM remodeling, therefore, their upregulation induced by AMG might contribute to better tissue remodeling during the wound healing.

**Fig. 4.**
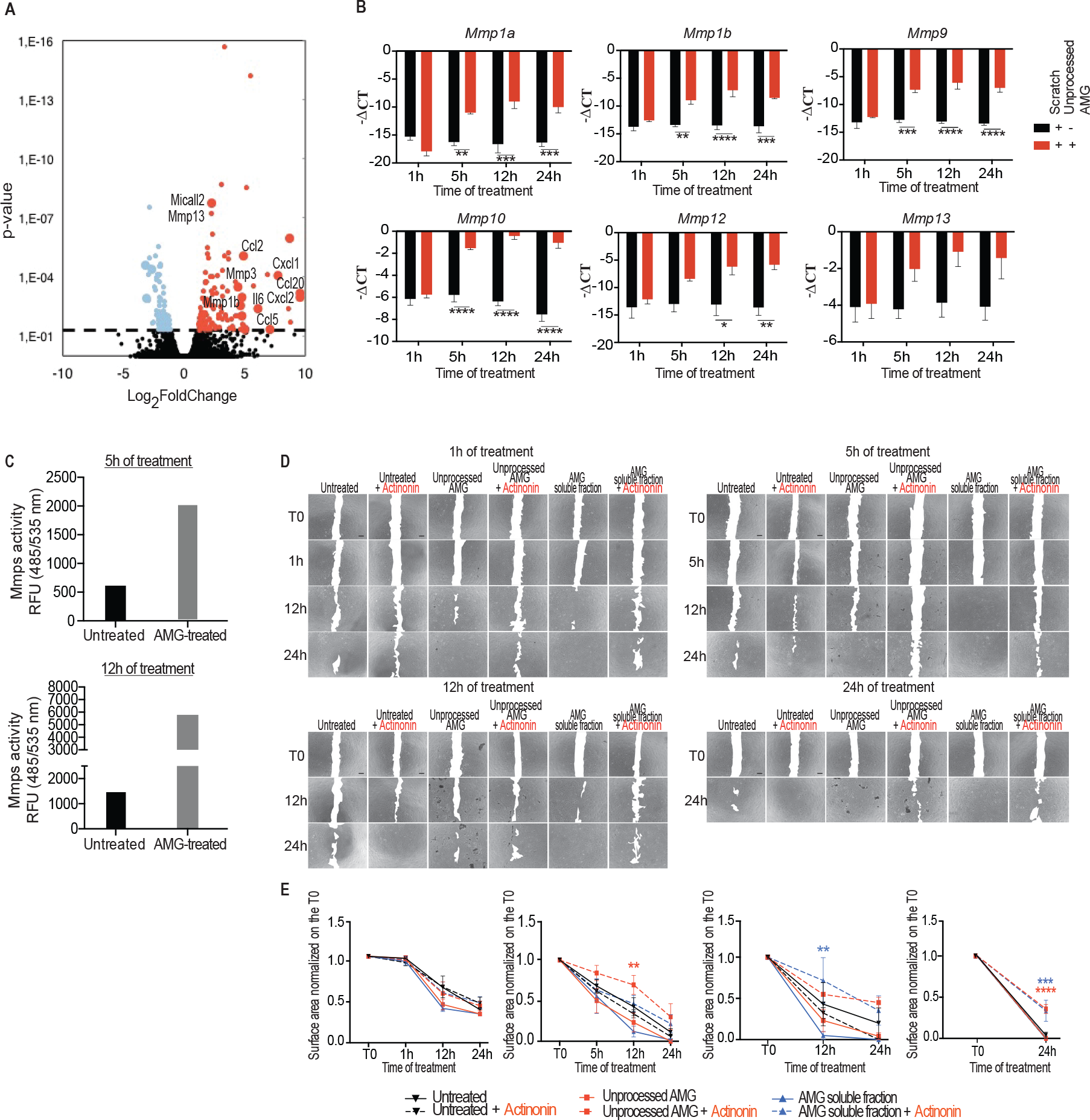
AMG-based treatment triggers matrix metalloproteinase expression and enzymatic activity in murine primary fibroblasts. **(A)** Volcano plot showing up- (red dots) and down-regulated (light blue dots) genes altered by the AMG treatment (5h of stimulation). Gene values are reported as a Log_2_FoldChange (p-value<0.05). **(B)** MMP gene expression evaluated by RT-qPCR. Expression values are expressed as a -ΔCT normalized on the expression of *Gapdh*, *B-actin* and *Rpl13a* housekeeping genes (3 biological replicates). Differences are calculated with two-way ANOVA (3 biological replicates) and indicated as * P < 0.05; ** P < 0.01; *** P < 0.005; ****P < 0.001. **(C)** Enzymatic activity of MMP members present in cell supernatant was fluorometrically detected. Data are presented as Relative Fluorescence Units (RFU). Signals were evaluated 30 minutes after starting the reaction using a microplate reader with a filter set of Ex/Em = 485/535. The fluorescence signal obtained from each sample was normalized on the substrate control. Groups of wounded cells that did not receive any treatment were used as a control. **(D)** Representative images of scratch wound assays on monolayers of murine primary fibroblasts. AMG was applied on wounded cells with different conditions (unprocessed AMG and its soluble fraction) and for different time periods (1, 5, 12 and 24h). Each condition was also incubated with the MMPs inhibitor – Actinonin – 20*μ*M for the same time periods of AMG treatment. Images were taken at the beginning (T0) and at regular intervals until wound closure was achieved. **(E)** Quantifications of AMG-treated cells with or without Actinonin are reported. Data are presented as mean ± SEM. Differences were calculated with two-way ANOVA (3 biological replicates) and indicated as ** P < 0.01; *** P < 0.005; ****P < 0.001.

Prompted by the results above, we investigated if increased MMP expression was accompanied by enhanced extracellular proteinase activity and accelerated cell migration. Upon scratch wounding and AMG exposition, we collected conditioned media and performed fluorescence-based assays of MMP enzymatic activity. AMG-treated cells showed a pronounced increase of MMP extracellular activity compared to the untreated condition (Fig. 4C). It therefore confirms our findings that fibroblasts increase MMP extracellular activity upon AMG treatment. Next, we set out to test whether the increase in MMP activity is responsible for higher cell motility. To that end we performed a scratch wound assay in cells treated or untreated with uprocessed AMG or enriched soluble AMG fraction in absence or presence of Actinonin for 1h, 5h, 12h or 24h. We first validated that the presence of Actinonin, a MMP-specific inhibitor, leads to significant reduction of MMP activity. (fig. S3A). Remarkably, scratch wound assays combined with inhibition of the MMP catalytic activity showed a decrease in the migration rate that led to a significant delay in scratch wound closure. These findings demonstrate that matrix MMP enzymatic activity plays a deterministic role in the AMG-dependent accelerated scratch wound closure (Fig. 4D and 4E). Of note, MMP expression is partially transcriptionally regulated in fibroblasts by AP-1 (*22*) a transcription factor formed by the dimerization of multiple components of the JUN, FOS, ATF and MAF protein families (*23*). Interestingly, we found that members of the AP-1 family such as *Fosl1*, *Fosl2*, *cJun* and *Junb* were highly up-regulated upon AMG treatment (fig. S3B). We also used publicly available ChIP-seq data to support that Fra1, a member of AP1 complex, binds to regulatory regions of MMPs (fig. S3C).

Taken together, our data demonstrate that AMG treatment increases MMP expression and extracellular activity, which are essential to mediate AMG-dependent accelerated cell migration.

### AMG treatment induces ERK–dependent MMP expression and cell migration

During WH, cellular functions are strictly orchestrated and regulated by the activation of several signaling pathways in order to efficiently achieve correct healing. GO and Network-based analyses associated with DEGs upon 5h AMG treatment showed enrichment of MAPK cascade and extracellular signal-regulated kinase (ERK1, ERK2 cascade; fig. S1H and S2D). In agreement, AMG treatment induced up-regulation of several target genes involved in ERK pathway regulation, as shown from the heat map (fig. S1I). In addition, all the growth factors mostly enriched in the AMG-solution, IGF-1, EGF and bFGF, are able to directly activate the intracellular ERK pathway (*24*).

On this basis we aimed to investigate whether AMG treatment affects the activity of the ERK pathway. We compared the levels of ERK phosphorylation (pERK) in scratched fibroblast monolayers untreated vs. AMG treated for 1h, 5h and 12h. We show that untreated fibroblasts increased pERK levels 5h after scratch injury with higher levels at 12h in agreement with the positive role of ERK pathway during cellular migration (*14*). In contrast, AMG treatment induced pERK at 1h with higher levels at 5h and 12h of treatment (Fig. 5A) supporting that AMG treatment accelerates ERK activation during the scratch wound assay. We next tested whether the ERK pathway is required for AMG-dependent acceleration of cell motility. To this end we performed scratch wound assays in fibroblasts treated or not with unprocessed AMG or enriched soluble AMG fraction in absence or presence of PD325901, a specific mitogen activated protein kinase kinase (MEK) inhibitor. Inhibition of the MAPK signaling pathway in either unprocessed AMG or soluble AMG fraction delayed significantly the capability of the cells to migrate compared to treatments without the MEK inhibitor (Fig. 5B and 5C). ERK inhibition was confirmed by western blotting analysis (fig. S4A). Furthermore, activation of the ERK signaling pathway has already been reported to correlate with increased expression of different members of the MMP family through AP-1 transcription factors binding (*25*). In accordance, we show that inhibiting MEK activity in AMG-treated fibroblast significantly reduced the levels of *Mmp1a*, *Mmp1b*, *Mmp9* and *Mmp10* gene expression (Fig. 5D). Surprisingly, levels of *Mmp12* were unchanged upon inhibition of MEK activity suggesting that *Mmp12* expression is regulated by AMG soluble factors but not by the MEK/ERK pathway (Fig. 5D).

**Fig. 5.**
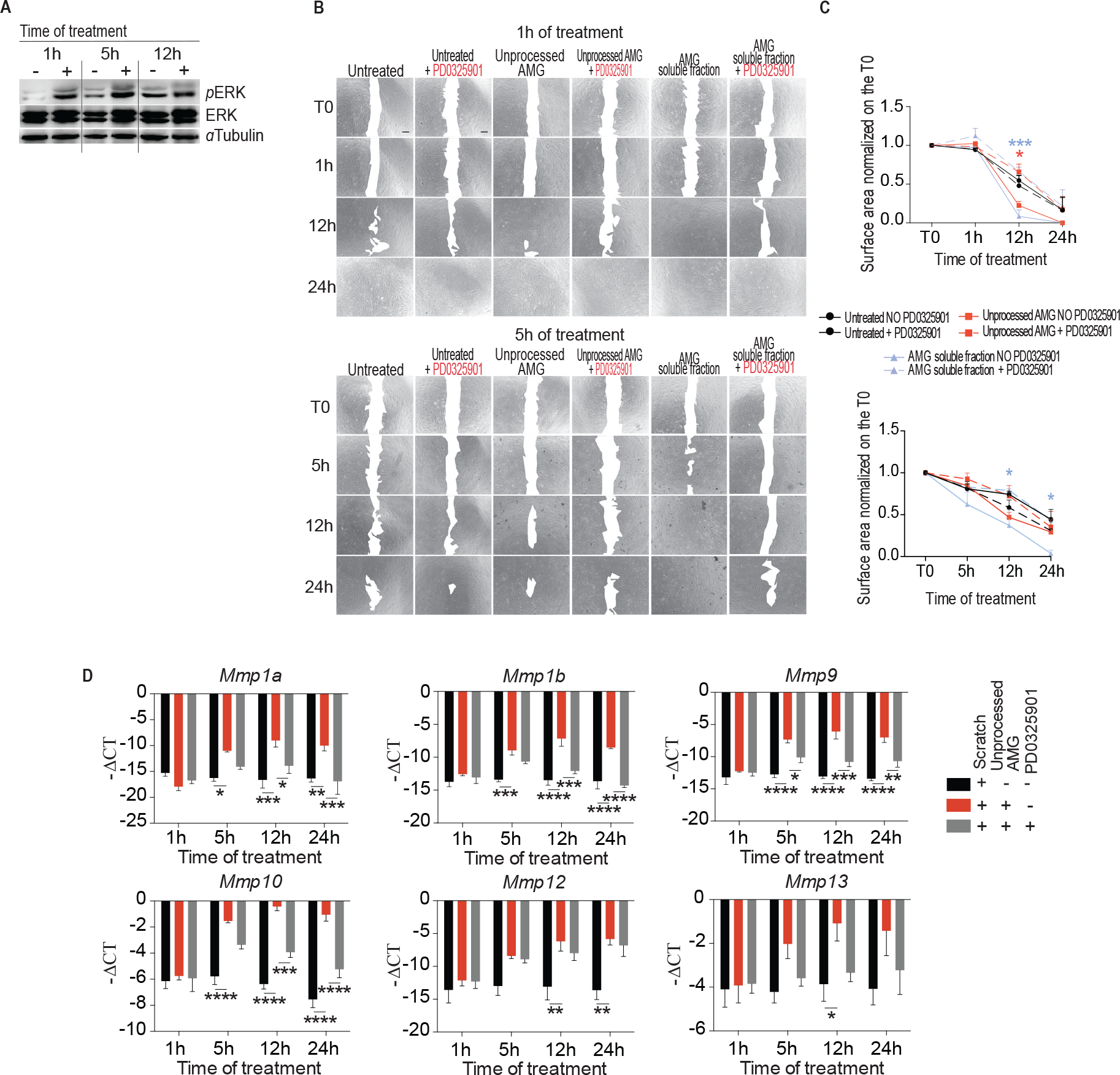
ERK/MAPK signaling pathway is specifically activated by AMG treatment. **(A)** Representative western blotting out of three biological replicates showing protein level of phosphorylated and total ERK1/ERK2 from wounded fibroblasts upon AMG treatment. **(B)** Representative images of scratch wound assays. Unprocessed AMG and its soluble fraction were applied on wounded cells. Each condition was also incubated with the MEK inhibitor – PD0325901– 1μM for the same time periods of the AMG treatment. Wounded cells did not receive any treatment and were used as a control. **(C)** Quantifications of each time of treatment were reported. Data are presented as mean ± SEM. Significant differences vs control were calculated with two-way ANOVA (3 biological replicates) and indicated as *P < 0.05 and ***P < 0.005. **(D)** MMP gene expression evaluated by RT-qPCR upon AMG treatment and MEK inhinitor PD0325901 exposition. Expression values are expressed as a -ΔCT normalized on the expression of *Gapdh*, *B-actin* and *Rpl13a* housekeeping genes (3 biological replicates). Differences are calculated with two-way ANOVA (3 biological replicates) and indicated as * P < 0.05; ** P < 0.01; *** P < 0.005; ****P < 0.001.

Collectively, our findings attribute the AMG-mediated WH to an increase in cell migration rate through ERK-dependent MMP activity, which may be regulated upstream by a pool of pro-motility growth factors.

### ERK signaling pathway drives autologous micrograft-mediated WH

Considering the strong AMG treatment ability to promote motility and migration of fibroblasts *in vitro*, we used the excisional WH mouse model to address whether topical application of AMG would also stimulate the healing capacity of full-thickness dermal wounds *in vivo*. In parallel, we tested the involvement of the MAPK cascade on *in vivo* AMG treatment by using the specific MEK inhibitor trametinib (*15*).

Full-thickness wounds were made on the back of C57BL/6 mice and splinted with silicone rings to limit wound closure caused by skin contraction. Subsequently, animal groups were topically and repeatedly treated on day 0, 2, 4, 6 and 8 with AMG (diluted in DMSO, 1:3), obtained from autologous skin biopsies, or vehicle control (PBS diluted in DMSO, 1:3), in the presence or absence of trametinib (Fig. 6A). We show that topical application of AMG significantly accelerated wound closure on days 4, 6 and 8, quantified as % of wound closure compared with mice treated with vehicle. Furthermore, topical administration of trametinib significantly delayed both vehicle and AMG-treated wound closure. This corroborates the essential role of the ERK signaling pathway in WH (Fig. 6B and 6C).

**Fig. 6.**
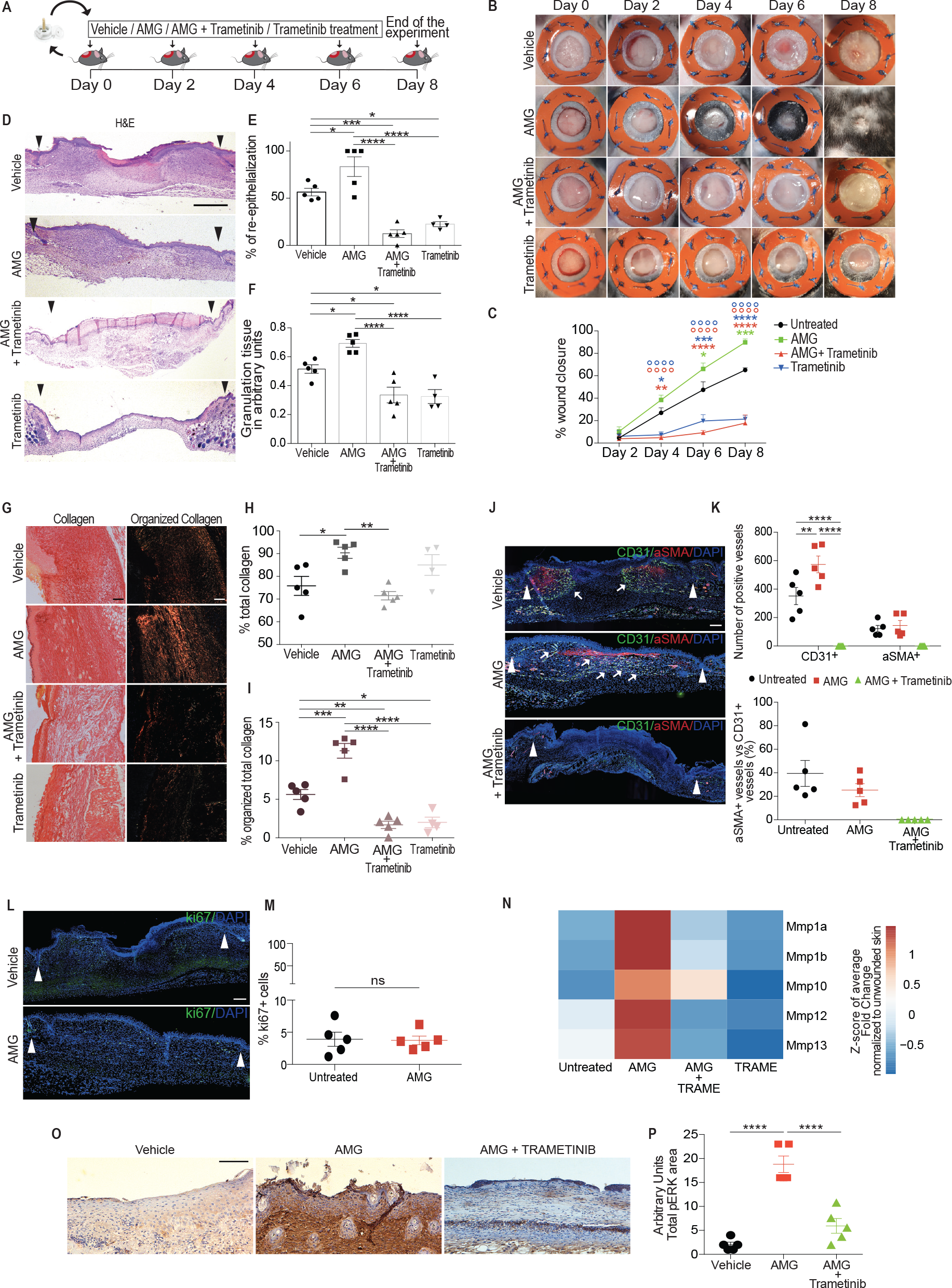
The *in vivo* healing potential of AMG through ERK signaling pathway. **(A)** Schematic representation of the *in vivo* WH study performed on C57BL/6 mice. Animals were topically treated with vehicle, AMG and MEK inhibitor – trametinib (0.2 mg) on day 0, 2, 4, 6 and 8. At the end of the experiment, skin wound samples were collected for further analyses. **(B)** Excisional wound-splinting assay showing the potential of AMG to improve wound closure in AMG-treated mice compared to the other conditions under study (vehicle, AMG + trametinib and trametinib). **(C)** Percentage of wound closure between the groups for each time point was calculated by the formula reported in the supplementary materials and methods **(D)** Representative hematoxylin and eosin (H&E) stained section on day 8 after wounding. Scale bar 500 μm. Arrowheads delimitate the wound area. **(E, F)** Percentage (%) of re-epithelialization and granulation tissue formation expressed in arbitrary units (AU) among the groups were evaluated. **(G)** Representative Sirius Red (SR) stained section on day 8 after wounding. Scale bar 100 μm. **(H, I)** % of total collagen formation and % organized collagen on the total collagen was evaluated. **(J)** Representative immunofluorescence of vehicle and AMG-treated wounds exposed or not to trametinib, showing presence of CD31+ and ɑ-SMA+ vessels. Scale bar 200 μm. Arrowheads delimitate the wound area. Arrows indicate CD31+ vessels. **(K)** Number of CD31+ and ɑ-SMA+ vessels and ɑ-SMA coated CD31+ vessels were evaluated in the entire wound area. **(L)** Representative immunofluorescence of vehicle and AMG-treated wounds showing the presence of the proliferative marker Ki67. **(M)** % of Ki67+ cells were evaluated (n=5). No significant differences were found. Scale bare 200 μm. Arrowheads delimitate the wound area. **(N)** MMP gene expression evaluated by RT-qPCR in vehicle and AMG-treated in presence or absence of trametinib. Expression values are expressed as a z-score of the average fold change (FC) normalized on the expression of *Gapdh*, *B-actin* and *Rpl13a* housekeeping genes (n=3). **(O)** Representative immunohistochemistry for pERK in vehicle and AMG-treated in presence or absence of trametinib. Scale bar 50 μm. All data are presented as mean ± SEM. Differences are calculated using two-way or with one-way ANOVA (n=5), corrected for multiple comparison and indicated as * P < 0.05; ** P < 0.01; *** P < 0.005; ****P < 0.001.

Next, we set out to evaluate whether AMG treatment positively affects a wide set of key WH parameters *in vivo.* First, we evaluated the benefit of AMG treatment on epidermal healing by analyzing the percentage of re-epithelialization. Among the overlapping and highly regulated sequential phases of the healing process, re-epithelialization plays a cardinal role regulating the closure of the injury. We found that none of vehicle-treated mice showed complete epithelial coverage of the wound at day 8 after damage, while wounds in three out of the five AMG-treated mice exhibited complete re-epithelialization. Re-epithelialization coverage in wounds treated with AMG reached 83% ± 10% compared to only 56% ± 4% in vehicle-treated mice (Fig. 6D and 6E). Importantly, mice treated with trametinib showed significantly reduced re-epithelialization compared to vehicle and AMG-treated mice (Fig. 6D and 6E). Next, we evaluated granulation tissue (GT) formation, new deposition and organization of collagen fibers as well as formation of new blood vessels. Fibroplasia and angiogenesis have both been shown to be indispensable events needed for forming new ECM and granulation tissue (*26*) and in turn successful WH. We show that the percentage of GT formation of AMG-treated group was significantly higher (70% ± 2,7%) compared to the other conditions, suggesting increased fibroplasia in AMG-treated mice (Fig. 6F). The total collagen content was statistically significantly greater in the AMG-treated wounds compared to the vehicle-treated as indicated by Sirius Red staining under bright light (Fig. 6G and 6H). Furthermore, the amount of organized (red birefringent) collagen, revealed with polarized light microscopy on the same cross-sections, was also significantly higher in AMG-treated compared to the control-treated wounds. Importantly, the presence of trametinib reduced both the percentage of polarized collagen and total collagen formation upon AMG treatment (Fig. 6I).

Immunofluorescence analysis of the endothelial marker CD31 as well as αSMA revealed that the AMG-treated group displayed increased levels of total CD31+ vessels but no significant differences were found in both number of total αSMA+ vessels and αSMA-coated CD31+ blood vessels compared to vehicle-treated mice, indicating the potential of AMG to trigger formation of blood vessels. However, as marked by low levels of αSMA the maturation of the newly formed vessels was still incomplete at the time of the analysis (Fig. 6J and 6K). Furthermore, in order to confirm *in vivo* that AMG treatment does not affect cell proliferation, wound areas were analyzed histologically and Ki67+ cells were quantified. No significant differences were shown between vehicle and AMG-treated mice in the % of Ki67+ cells (Fig. 6L and 6M).

Finally, MMP expression levels were analyzed 8 days after damage by RT-PCR in all the groups under study. Interestingly, levels of *Mmp1a*, *Mmp1b*, *Mmp10*, *Mmp12* and *Mmp13* were increased in AMG-treated mice. Treatment with the MEK/ERK inhibitor trametinib reduced expression of all five MMPs *in vivo* suggesting that MMP expression during *in vivo* AMG-mediated accelerated WH is ERK activation-dependent (Fig. 6N). In addition, phosphorylation of ERK was evaluated by immunohistochemistry, confirming the potential of AMG treatment to trigger the cascade (Fig. 6O and 6P).

Altogether, our results demonstrate the ability of AMG treatment to actively trigger all wound healing phases leading to an acceleration of the entire process.

## Discussion

Autologous micrograft (AMG) technique has gained attention in the last years for wound healing (WH) application. The potential of this technique has been shown to provide positive clinical outcome with significant WH acceleration (*2, 21*). However, the precise molecular mechanism through which AMG exhibits its beneficial effects remains unclear. In efforts to identify the molecular mechanism that drives the AMG beneficial effect in WH, we analyzed the gene expression signature of untreated and AMG-treated wounded murine primary fibroblasts. Using RNA sequencing followed by PPI network analysis we identified the AMG-specific WH signature, characterized by enrichment of important WH-related biological processes and signaling pathways, including cell migration, angiogenesis and MAPK/ERK cascade activation. Our work revealed that AMG therapeutic effect in both *in vitro* scratch closure as well as during *in vivo* WH assay induces an increase in cell migration rate, leading to an accelerated WH without affecting cell proliferation. Most importantly, we coupled this effect to increased MMP enzymatic activity sustained by the ERK signaling pathway. In accordance with our *in vitro* results, topical application of the specific MEK inhibitor – trametinib – impedes *in vivo* AMG-beneficial effects. Altogether, we provide a framework for how AMG supports the WH process opening avenue for further advances.

Upon injury, the earliest event that drives the progress of the WH response is characterized by gradients of cytokines/growth factors, such as interleukin-1 (IL-1), interleukin-6 (IL-6), EGF, PDGF, TGF-beta and bFGF, which will activate intracellular signaling pathways to encourage fibroblast migration towards the wound site. In the past years, several bio-active therapeutic approaches and cell-based therapies have been implemented to improve the healing process, including the delivery of growth factors or cells into a damaged tissue in order to improve the healing process (*27, 28*). Unfortunately, clinical trials of single growth factor-delivery treatment resulted in poor outcomes (*4*), suggesting that replacement of a single factor is not sufficient to improve the WH process. Rather, a combinatory treatment composed of pool of growth factors may provide a way to actively improve therapeutic WH approaches (*27*).

Previous studies have suggested that the presence of Mesenchymal Stem Cells (MSCs) within the AMG may drive their positive outcome in WH (*2, 21*). In this work and for the first time, we can mainly exclude the possibility that this is the main mechanism of action of the AMG treatment, as the AMG soluble fraction showed higher pro-motility effects on fibroblast than unprocessed AMG. Moreover, in accordance with the role of growth factors as one of the most important events to initiate the WH response, we showed that AMG solution is enriched in several positive regulators of cell motility, such as EGF, PDGFa, bFGF, IGF-I and IGFBPs (Figure 3D). In addition, AMG treatment on fibroblasts stimulates chemokine expression such as *Cxcl1*, *Cxcl2* and *Ccl2*, which play essential roles in immune cell chemotaxis and during the re-epithelialization phase (Fig. 4A). In agreement, previous studies showed that lack of these signaling molecules leads to *in vivo* WH delay with detrimental effects on both re-epithelialization and angiogenesis phases (*29*). In this frame, we show that AMG-based treatment advances the starting signal of the healing process, through the delivery of a pool of WH-associated growth factors that promote fibroblast migration and faster wound closure.

The growth factors present in AMG are ligands of receptor tyrosine kinases, the stimulation of which initiate activation of several biological processes and intracellular signaling pathways. To investigate active phenomena associated with the AMG-mediated WH, we analyzed enriched gene ontology (GO) biological processes and signaling pathways of AMG-associated DEGs.

Specifically, transcriptome analysis of AMG-treated fibroblasts clearly showed upregulation of the MAPK/ERK-signaling pathway, suggesting its involvement in the AMG molecular mechanism. It has been previously shown that activation of the ERK signaling cascade is positively involved in WH mainly through its capacity to stimulate cell proliferation (*15*). On the other hand, recent evidences show the role of the MAPK pathway in the enhancement of cell migration and morphology, angiogenesis (*30*) and, most importantly, in regulating tissue repair functions in fibroblasts (*31*). In line with these findings, our study demonstrates that AMG-dependent ERK activation leads to stimulation of migrating fibroblasts without effect on cell proliferation in the AMG-mediated accelerated WH (fig. S5). Moreover, AMG-treated wounds confirmed ERK-dependent AMG beneficial effects, showing improvement in several WH events, such as reepithelialization, granulation tissue formation and angiogenesis. Conversely, topical application of the specific MEK inhibitor - trametinib – caused significantly reduced wound closure, reepithelialization, granulation tissue formation, collagen deposition and organization as well as angiogenesis. These results highlighted the essential role of the ERK signaling pathway activation in the AMG-mediated WH. Very likely many signaling pathways might be involved in the AMG molecular mechanisms, including the TGF-beta signaling pathway (*32*). However, we show that inhibition of ERK pathway activation in AMG-treated wounds leads to an arrest of the WH progress. These findings suggest that AMG treatment was not able to exert its beneficial effects in the presence of the MEK inhibitor, confirming that ERK signaling activation is highly required for AMG-mediated WH.

In this study we show that AMG treatment increases MMP gene expression and their enzymatic activity (Fig. 4B; 4C). These family of proteases play pivotal roles in several biological processes involving proteolysis, including developmental processes (*33*), inflammation and wound healing (*34*) Our work revealed a specific WH-associated MMPs transcriptional signature in AMG-treated fibroblasts, where *Mmp1a*, *Mmp1b*, *Mmp9*, *Mmp10*, *Mmp12* and *Mmp13* are highly up-regulated as well as enzymatically activated upon AMG (Fig. 4A, 4B and 4C). Moreover, we show that activation of these proteases is coupled with an acceleration of migration rate in our *in vitro* WH model (Fig. 4D and 4E) in line with other studies (*6*). Remarkable, it has been previously shown that MMP-mutant mice exhibit impaired WH conditions with reduced level of cell migration (*6*). Mechanistically, we reveal that inhibition of the MMPs enzymatic activity, obtained using the specific MMPs inhibitor – actinonin – reduced both *in vitro* scratch closure and enzymatic activity, demonstrating their important role in the AMG molecular mechanism. MMPs expression can be specifically stimulated by cytokines and growth factors, including EGF, FGF and many others, which in turn regulate AP-1 family members activity through MAPK signaling pathway (*25*).

Interestingly, we show that the promoter regions of all the up-regulated MMPs contain binding sites for the AP-1 transcription factor, which activation is known to be sustained by ERK signaling pathway (*35*) (fig. S5). Here, our results show AMG-dependent upregulation of AP-1 family members including *Fosl1*, *Fosl2*, *cJun* and *Junb,* the activation of which was reverted upon trametinib (fig. S4B). Similarly, our work demonstrates that MMPs gene expression regulation specifically depends on ERK activation, being down-regulated upon MEK inhibitor (Fig. 5D).

However, *Mmp12* expression, which was also significantly up-regulated upon AMG, showed no differential expression after MEK/ERK inhibition, suggesting that other signalling pathways might be involved in the AMG-mediated *Mmp12* regulation. All in all, our study highlights the potential of AMG treatment in accelerating the healing process through increased cell migration obtained by an ERK-dependent MMPs regulation. We, therefore, speculate that ERK-mediated MMP stimulation observed upon AMG treatment may depend on AP-1 transcription factor regulation.

Taken together, our study demonstrates the inherent ability of AMG treatment in promoting the endogenous regenerative capacity and provides insights into the AMG molecular mechanism proving its ability to trigger the whole WH process. Therefore, we offer a framework for future improvements of autologous micrograft therapy for skin tissue regeneration.

## Materials and Methods

### Study design

All the experiments performed in this study were approved by the Ethics Committee at KU Leuven university (ethical approval codes P056/2017 and P036/2018). Primary adult fibroblasts were obtained from murine tail’s tips and cultured until full confluency. The scratch wound assay was then performed using a pipette tip, creating an artificial wound on cell monolayers (*36*). On the other hand, Autologous Micrograft (AMG) solutions were obtained starting from skin biopsies (1 cm^2^), through a method of disintegration using a sterile tissue destroyer (*32*). 1 mL of AMG solution was diluted 1:10 in growth media and applied on an *in vitro* scratch wound assay. Inhibitory drugs for MEK (PD0325901 1 *μ*M) and MMP activity (Actinonin 20) *μ*M were applied on scratched monolayers. Images were taken at the beginning (T0) and at regular intervals until scratch wound closure was achieved. Images were then analyzed with Image J and the surface areas were evaluated.

RNA samples of murine wounded fibroblasts exposed to different times of AMG treatment (1, 5, 12 and 24 hours) were collected, sequenced and processed by the Genomics Core (KU Leuven – UZ Leuven, Belgium).

*In vivo* WH was evaluated by performing the excisional wound splinting mouse model (*37, 38*) (supplementary materials – materials and methods). Specifically, four groups of C57BL/6 mice (12 weeks old, n=5 per group) were anesthetized, shaved and excisional wounds were made in the shaved dorsal skin under ketamine/xylazine anesthesia with a 5-mm biopsy punch. Silicon rings were sutured around wounds in order to limit wound closure caused by skin contraction. Wounds were then topically treated with thirty microliters of AMG (diluted in DMSO, 1:3), obtained from the 0.5 cm of murine skin biopsy (*32*). A second group (n=5) was topically treated with AMG supplemented with a specific MEK inhibitor – trametinib (0.2 mg, Cat No. HY-10999, MCE). Trametinib was dissolved in dimethylsulfoxide (DMSO) to a concentration of 20 mg/ml and 10 *μ*L of the mixture (corresponding to 0.2 mg) was added topically together with the AMG. A third group (n=5) was topically treated with the MEK inhibitor – trametinib (0.2 mg, Cat No. HY-10999, MCE) (1:3 in PBS). The control group was treated with PBS diluted with DMSO 1:3 as vehicle. Subsequently, wounds were covered with Tegaderm™ dressing (3M, Maplewood, USA) to protect the wound area and to prevent the area from drying out. Every other day (day 2,4, 6 and 8), digital images were taken using a Canon EOS 5D Mark II camera, treatments were re-applied topically to the wound site and dressings were renewed. Changes in percentage of wound closures were calculated by comparing the healed wound area of specific days with the original wound area over the time (*38, 39*) (supplementary materials – materials and methods). At day 8, animals were sacrificed and dorsal skin wound samples were collected for further histological and gene expression analyses. Several WH parameters were evaluated during the study, including % of reepithelialization, granulation tissue formation, collagen synthesis and organization, angiogenesis and cell proliferation. Hematoxylin and eosin, Sirius Red, CD31/alpha-SMA, Ki67, p-ERK immunostaining as well as MMP gene expression analyses were performed on the skin samples unveiling the AMG-mediated effect on the different groups under study.

### Statistical Analysis

For RNA sequencing data, statistical analysis was performed using bioconductor package DESeq. Reported p-values were adjusted for multiple testing with the Benjamini-Hochberg procedure, which controls false discovery rate (FDR). GraphPad Prism 6 was used to graph the data and GraphPad Prism software (San Diego, CA, USA) was used to analyze all the experiments of this study. Significant differences were determined by one-way, two-way analysis of variance (ANOVA), adjusted for multiple comparison and two-tailed unpaired t test. Data are presented as mean ± standard error of mean (SEM) or as fold change.

## Acknowledgments

We thank to Hanne Grosemans and Petra Vandervoort for technical support.

### Funding

This work is supported by the contribution of NATO grant “RAWINTS” project G984961: Rapid Skin Wound Healing by INtegrated Tissue engineering and Sensing. We are grateful for the support from KU Leuven starting grant (STG), KU Leuven C1 funds (C14/16/078) and FWO (G097618N) funds to F.LL; FWO (#G088715N, #G060612N, #G0A8813N). CARIPLO Foundation #2015, C1 funds (C14/17/111) and Opening the future #EJJ-C4851-17/07-P to M.S.

### Author contributions

M.B., F.L.L. and M.S. conceived, designed and performed all the *in vitro/in vivo* experiments and bioinformatics analysis, wrote and revised the paper. F.V designed and developed the bioinformatics pipeline, wrote and revised the paper. A.C-C. contributed to the bioinformatics analysis. A.J. contributed to the bioinformatics analysis, data interpretations and revised the paper. E.C. and A.L. contributed to the *in vivo* experiments and revised the paper. R.D. contributed to the FACS data analysis, F.R., G.C., M.G.C.D.A. and R.B. revised the paper.

### Competing interests

All authors declare that they have no competing interests.

### Data and materials availability

RNA sequencing data are deposited in GEO Datasets under accession number GSE123829.

## Supplementary materials

### Materials and Methods

#### Cell culture

Mouse primary fibroblasts were isolated from the end of the tail (0.5 cm) of C57BL/6 mouse model under anesthesia (isoflurane). Tails were disaggregated by collagenase IV digestion for 40 minutes at 37°C in agitation. The obtained cells were cultured in DMEM GlutaMax (Ca No. 61965-026, Gibco Life Science, Grand island) supplemented with 10% of fetal bovine serum, 1% of penicillin-streptomycin, 1% of L-glutamine, 1% of sodium pyruvate and 1% of nonessential amino acids at 37°C in a humidified incubator with 5% of CO_2_ and maintained in culture until obtaining the correct level of confluence (90%) to perform the cell scratch assay.

#### Preparation of Autologous micrografts

Autologous micrografts (AMG) were obtained from skin murine biopsies (1 cm^2^, collected from the back of the animals), using a sterile tissue destroyer (Rigeneracons*®*)(*32*). Tissues were re-suspended in 1 mL PBS, disrupted, and filtered through 70 *μ*m filter membranes. The extracted AMG were then diluted 1:10 in growth media and applied on murine primary fibroblasts on an *in vitro* model of wound healing (cell scratch assay).

#### Cell scratch assay

Mouse primary fibroblasts were plated at a density of 3500 cell/cm^2^ on 24 multi-well culture plates. Cells at passages 4 to 6 were used in this study. After reaching confluence, a scratch was performed on the monolayer of cells using a pipette tip, resembling an artificial *in vitro* wound. The *in vitro* cell scratch assay represents a cost-effective technique to measure cell migration *in vitro*. After the creation of the damage, debris were washed out using sterile PBS (phosphate-buffered saline) and images were taken at the beginning (T0) and at regular intervals until the closure of the *in vitro* scratch was achieved. The comparison between the images allowed quantifying the cell migration rate. AMG treatment was applied on the wounded cells for different time of treatment (1h, 5h, 12h and 24h).

#### XTT cell viability assay

The *in vitro* cell viability was assessed evaluating metabolic activity and micrograft-related sensibility in primary murine fibroblasts exposed to the AMG, following manufacturer instructions (purchased from Thermo Fisher Scientific). In brief, mouse primary fibroblasts were plated at a density of 10^5^cell/well, in 96 multi-well culture plate, in 100 µL of Growth medium, in which AMGs were diluted (1:10) followed by different time period of treatment. At the end of each time period, cell culture media was replaced with 100µL of XTT/PMS (Phenazine methosulfate) (1 mg/ml; 10 mM) solution. Cells were incubated for 2 hours at 37°C. The absorbances were read at 450 nm by Nanodrop 1000 spectrophotometer (Thermo Fisher Scientific). Absorbance values are normalized to the control (cells who not received the AMG treatment), plotted using GraphPad Prism 6 and presented as FoldChange.

#### Cell cycle assay

Changes in the cell cycle were assessed using the EdU - 5-ethynyl-2’-deoxyuridine assay (Thermo Fisher Scientific). Scratched cells were treated with the AMG (1h, 5h, 12h and 24h of treatment) or with vehicle (PBS). After each time point, cells were incubated at 37°C with EdU 10 *μ*M for 2 hours. Cells were harvested with Trypsin-EDTA 0,25% (Ca No. 25200056, Gibco, Grand island, NY, USA), washed with PBS 2% FBS and pelleted. After washes and permeabilization steps, (Samples were fixed with 4% PFA for 20 minutes at RT, washed with PBS 2% FBS, permeabilized with Triton-X100 0,5% in PBS for 20 minutes at RT and washed again with PBS 2% FBS) cells were incubated with 250 *μ*L of cocktail solution (containing PBS 1X, CuSO_4_ 100 mM, Sodium Ascorbate 1M, Azide Alexa Fluor 647) for 10 minutes at RT in the dark, followed by two washes with PBS 2% FBS and pelleted. 500 *μ*L of PBS 2% FBS was finally added on the cells supplemented with 1 *μ*L of Propidium Iodide (PI) and RNase 250 *μ*g/mL for 30 minutes at RT. The samples were then acquired by Flow Cytometry (BD-Biosciences) and analyzed by FlowJo software.

#### Immunofluorescence staining

20^5^/well of Murine primary fibroblasts were plated at a density of on 24 multi-well culture plates. After reaching a confluent cell layer, scratch assay was perform using a 200 *μ*L pipette tip. Cells were fixed with 4% PFA for 20 minutes at RT, permeabilized with 1% BSA, 0,2% Triton-X in PBS for 30 minutes at RT, washed repeatedly and blocked with donkey serum 1:10 in PBS (blocking solution) for 40 minutes at RT. On the other hand, wound murine biopsies were collected and further processed for paraffin embedding. Cells and 7 *μ*M wound skin sections were incubated overnight at 4°C with primary antibodies. Primary antibodies used in this study were Mouse monoclonal anti-actin α-Smooth Muscle - Cy3™ (1:200, C6198, Sigma Aldrich), The following day, after PBS washes, the cells were incubated with secondary-AlexaFluor antibody for 1h at RT and washed repeatedly with PBS. Hoechst 33342 (1:10000) was used for the nuclear counterstaining, samples were washed three times with PBS and wells/slides with cells were mounted using FluorSave™ mounting medium (EDM Millipore). The images were taken using Nikon Eclipse Ti Microscope and NIS-Elements AR 4.11 software. Pictures were analyzed, quantified and merged using ImageJ software (NIH).

#### Western Blotting

Scratched murine fibroblasts treated with AMG for 1h, 5h, 12h and 24h or untreated cells were lysate in RIPA Buffer containing 0,5 mM Sodium Orthovanadate, 1:100 Protease inhibitor cocktail and 1mM Phenylmethanesulfonyl Fluoride. Equal amount of protein extracted from each sample (30 *μ*g) was heat-denaturated in sample-loading buffer (50 mM of Tris-HCl, pH 6.8, 100mM DTT, 2% SDS, 0.1% Bromophenol Blue, 10% Glycerol), resolved by SDS-PAGE and transferred to nitrocellulose membranes (GE Healthcare Bio-Sciences, Pittsburgh, USA). Membranes were then stained with Ponceau S dye (Sigma-Aldrich) and were blocked with non-fat dry milk 5% in TBS (Tris-Buffered Saline) supplemented with Tween 0.05% for 1 hour at RT. Membranes were then incubated overnight at 4°C in agitation with primary antibodies: rabbit α-Smooth Muscle actin (1:800, ab15734, Abcam), mouse p-ERK (1:1000, sc-7383, Santa Cruz), rabbit ERK (1:1000, sc-94, Santa Cruz), mouse Tubulin-alpha (1:1000, T5168, Sigma Aldrich). The following day, membranes were incubated with secondary horseradish peroxidase HRP-conjugated antibodies (1:5000, Santa Cruz Biotechnology, CA, USA), in TBS 0.05% tween and non-fat dry milk 2,5%. Analyses were performed using the ChemiDoc XRS+ detection system (BioRad, Temse, Belgium) in combination with chemiluminescent HRP substrates SuperSignal West Pico PLUS and SuperSignal West Femto (Thermo Fisher Scientific). Relative densitometry was obtained normalizing all samples to the housekeeping Tubulin-alpha, using both the QuantityOne software and ImageLab software (BioRad).

#### Growth factors antibody array

Simultaneous detection of several growth factors within the AMG was assessed using Mouse Growth Factor Antibody Array Membrane containing 30 targets (RayBiotech, AAM-GF-3-2). The membranes (spotted with 30 different antibodies) were blocked using the provided blocking buffer for 30 minutes at RT, then incubated with AMG for an equal amount of protein 30 *μ*g (1:10 diluted in blocking buffer) overnight at 4°C. The following day, membranes were overnight incubated at 4°C with provided Biotinylated Antibody Cocktail. After the washes step, the membranes were incubated with HRP-conjugated Streptavidin (1X) for 2 hours at RT and washed again. Chemiluminescent detection was obtained using the provided detection buffers. Relative densitometry was obtained using the QuantityOne software (BioRad) and normalizing all signals to the positive controls signals of each membrane.

#### Matrix metalloproteinases (MMPs) enzymatic activity

MMPs activity was detected using MMP activity Assay Kit ab112146 (Abcam), following manufacturer instructions. The enzymatic activity of all MMPs members present in cell supernatant was detected fluorometrically. Data are presented as RFU (Relative Fluorescence Units). Signals were evaluated 30 minutes after starting the reaction using a microplate reader with a filter set of Ex/Em = 485/535. The fluorescence signal obtained from each sample was normalized on the substrate control.

#### Transwell Migration Assay

Murine primary fibroblasts were detached using 0,25% Trypsin-EDTA, washed with PBS (1X) and pelleted. Transwell Multiple Well Plate with Permeable Polycarbonate Membrane Inserts (Cat No. 3422, Thermo Fisher Scientific) were used for this study. 10^5^cell were plated on the upper layer of the transwell in a maximum volume of 100 *μ*L of cell culture media. 600 *μ*L of unprocessed AMG (1:10 diluted in Growth medium) or 600 *μ*L of AMG soluble fraction (1:10 diluted) were added in the lower part of the chamber. A well containing 10^5^cell and 600 *μ*L of complete GlutaMax media were used as a control. After incubation of 24 hours, the transwell inserts were removed from the chemo-attractant factors, cleaned from remaining media and not migrated cells using a cotton-tipped applicator and fixed with 70% ethanol. After the fixation time, membranes were left to dry and stained with 0,1% of crystal violet. After the staining time, membranes were detached from the inserts, deposited on slides and mounted using DPX. Picture of the membranes were taken with Axiovert 40 CFL connected with Axiocam MRc5 (Zeiss) and analyzed using ImageJ software.

#### RNA extraction, mRNA gene expression

Total RNA was purified using PureLink^®^ RNA Mini Kit (Cat No. 12183018A, Life Technologies™) according to the manufacturer’s instructions. cDNA was generated using 0,5 *μ*g of RNA and obtained by the Superscript III Reverse Transcriptase First-Strand Synthesis SuperMix (Invitrogen). Quantitative real-time PCRs (qRT-PCR) were performed using the Platinum^®^ SYBR^®^ Green qRT-PCR SuperMix-UDG (Cat No. 11733038, Thermo Fisher Scientific) on a ViiA™ 7 Real-Time PCR System with 384-well plate (Cat No. 4453536, Applied Biosystems). Gene expression values were normalized based on the *Gapdh*, *B actin*, *Rpl13a* housekeeping genes. All primers that were used were purchased from IDT technologies, Leuven, Belgium and are reported in table S4.

*RNA sequencing and bioinformatics analyses* (deposited in GEO Datasets under accession no. GSE123829). RNA samples were, quantified with Nanodrop 1000 spectrophotometer (Thermo Fisher Scientific) and RNA integrity was evaluated using Bioanalyzer (Agilent 2100) combined with Agilent RNA 6000 Nano Kit (Ca No. 5067-1511). RNA samples were then processed by the Genomics Core (KU Leuven - UZ Leuven, Belgium). Library preparation was performed with the Illumina TruSeq Stranded mRNA Sample Preparation Kit, according to the manufacturers protocol (48 samples, Cat No. 20020594). Denaturation of RNA was performed at 65°C in a thermocycler and cooled down to 4°C. Samples were indexed to allow for multiplexing. Sequencing libraries were quantified using the Qubit fluorometer (Thermo Fisher Scientific, Massachusetts, USA). Library quality and size range was assessed using the Bioanalyzer (Agilent Technologies) with the DNA 1000 kit (Agilent Technologies, California, USA) according to the manufacturer’s instructions. Libraries were diluted to a final concentration of 2nM and sequenced on Illumina HiSeq4000 sequencing system, according to the manufacturer’s recommendations. 50 bp single-end reads were generated. An average of 20 million reads were obtained from all samples. Quality control of raw reads was performed with the FastQC v0.11.5. The adapters were filtered with ea-utils v1.2.2.18. The splice-aware alignment was performed with TopHat v2.0.13 against the mouse genome mm10. The number of allowed mismatches was 2. Reads that mapped to more than one site to the reference genome were discarded. The minimal score of alignment quality to be included in count analysis was 10. Resulting SAM and BAM alignment files were handled with Samtools v0.1.19.24. Quantification of reads per gene was performed with HT-Seq count v0.5.3p3. Count-based differential expression analysis was done with R-based (The R Foundation for Statistical Computing, Vienna, Austria) Bioconductor package DESeq. Reported p-values were adjusted for multiple testing with the Benjamini-Hochberg procedure, which controls false discovery rate (FDR).

#### N-of-1 pathway MixEnrich single-subject analysis (SSAs)

The N-of-1 pathway MixEnrich is an approach designed to analyse gene expression data from paired samples drawn from the same subject when large cohorts or replicated are not available (*18*). An overview of this approach is reported in Fig. 1SA. We applied this technique to the paired samples (AMG-treated vs untreated cells) drawn from the same animal upon different AMG treatment time points (1h, 5h, 12h, and 24h). This produced a total of 8 paired samples. All samples were first filtered normalized by using NOIseq (*40*), since it is considered more reliable with SSA (*41*). Next, for each transcriptome sample the absolute value of log transformed fold change *-|log_2_FC|* (Fig. 1SA) was obtained as *|log_2_(U/T)|* where *U* is the expression level of a gene in the untreated condition and *T* is its expression level in the AMG-treated condition. Genes were then clustered in dysregulated genes (*cluster a*) and non-dysregulated (unaltered) genes (*cluster b*), based on the *-|log_2_FC|* values (Fig. 1SA). The clusters members have been assigned using a Bernoulli trial (Eq.1)

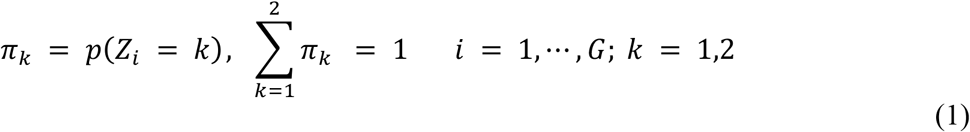

where *Zi* the latent variable and *G* is the total number of genes detected in the transcriptome. The clusters formation was then followed by evaluation of the probability that each gene belongs to the assigned cluster, evaluated by posterior probability by using Bayes rule (Eq.2)

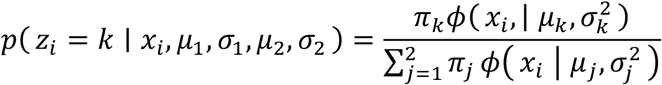

Dysregulated genes have been defined by a posterior probability above 0.5, where this cluster is defined as the cluster with the larger mean. After allocating the mRNAs to specific clusters, the enriched pathways, related to each time period of treatment were obtained using a Fisher’s Exact Test (FET). The FET is computed according with the contingency table (table S1), assuming one pathway consists of *M* genes among which *d* genes are dysregulated; while the entire genome consists of *N* genes among which *D* genes are dysregulated. Thanks to this procedure, pathway dysregulation is determined relative to the dysregulation of the entire transcriptome as the background for each AMG time of treatment (1h, 5h, 12h and 24h) (table S1). Since different pathways may not be independent due to overlapping genes between them, the p-values resulting from FETs were adjusted for multiple hypothesis testing using Benjamini and Yekutieli approach that accounts for correlated p-values.

#### Differential expression and enrichment analysis

List of differentially expressed genes, obtained from the our expanded cohort of samples (n=3 biological replicates), were selected at a adjust p-value < 0.05 and used to perform enrichment analysis, through Gene Ontology (GO) via Panther classification system (table S2), as well as used to build the Protein-Protein Interaction (PPI) network representing the wound healing process, by using the public PPI repository STRING. Enriched biological processes and pathways were selected at FDR < 0.05 and presented using GraphPad Prism 6. Among these lists, selected GO terms were then used to obtained either specific cell migration/cell motility signature genes or ERK/MAPK signature genes, which were then presented as heat maps (via R studio software).

#### Network construction

The PPI network was constructed using the repository STRING v.10.5 (*20*). We downloaded from STRING all the PPIs related to the mouse organism and, according with our previous works (*42, 43*), we retained the PPIs based on database or experimental evidences and with STRING confidence score higher than 700. These interactions are considered the most reliable, for more details please refer to (*42, 43*). We constructed the network starting from the protein codified by the DEGs resulting from the previous RNA-Seq data related to AMG treatment at 5 hours. We used these proteins as seed nodes and we linked them to the related STRING interactors. We finally checked if additional PPIs were present in STRING between all the added nodes and included them in the network as edges. Note that the constructed network is also a weighted network where the edge weights correspond to the STRING confidence score associated to the PPI (edge).

#### Hub nodes

We identified the network hubs by retaining the top 10% of the highest degree nodes, i.e. number of neighbors. This threshold was suggested by other studies (*42*). This allowed to identify nodes having a key role in the network and therefore in the AMG treatment process. Several studies in fact demonstrated that hubs likely correspond to network nodes playing an important role in the system represented (table S3) (*44, 45*).

#### Network Clustering

It is known that topological clusters corresponds specific biological functions or biological processes (*46*). Therefore, we performed the network clustering and subsequently, the biological process enrichment of the nodes in each cluster to study the behaviors of gene groups clustered together based on their topological properties. In our study, we performed the clustering using the Cytoscape (*47*) plugin ClusterONE (*48*) because of its reliability and since it considers network weights (i.e., STRING confidence scores) and allows cluster overlapping (*48*). Statistically significant clusters are identified by a *p*-value, also provide by ClusterONE through a one-sided Mann-Whitney U test performed on the in-weights and out-weights of cluster vertices. A low *p*-value indicates that the in-weights are highly larger than out-weights, leading to a reliable cluster which is not the result of random fluctuations. To distinguish significant clusters from not statistically significant ones, authors suggest a *p*-value threshold of 0.05 (*48*) (table S3).

#### Hub and clusters enrichment analyses

We performed the enrichment of a group of gene, such as hubs or genes belonging to a clusters obtained using the Cytoscape plugin ClueGO (*49*). ClueGO allows to directly load gene group of interest from a network, to perform the enrichment analysis using different source of knowledge (e.g. GO), to extract the list of significant terms and to obtain a network representation of the biological processes resulting significant. Here, we applied this method to hub nodes and to each cluster group of nodes. We considered enriched all the GO-BPs whose p-value was smaller than 0.05 by setting directly in ClueGO Cytoscape plugin this threshold.

#### ChIP-sequencing analysis

Publicly available ChIP-seq data (*50*) was processed using the pipeline from the Kundaje lab (Version 0.3.3). Reads were aligned to reference genome (mm10) using Bowtie2 (v2.2.6) using the ‘--local’ parameter. Single-end reads that aligned to the genome with mapping quality ≥30 were kept as usable reads (reads aligned to the mitochondrial genome were removed) using SAMtools (v1.2). PCR duplicates were removed using Picard’s MarkDuplicates (Picard v1.126). Coverage was calculated using bamCoverage function with binsize = 1 bp and normalized using RPKM.

#### In vivo WH assay

20 12 week-old female C57BL/6 mice were used in this study. Excisional wound splinting mouse model was performed (*37, 38*). Mice were shaved, disinfected with betadine and ethanol and full-thickness wounds (0.5 cm diameter) were made in the shaved dorsal skin under ketamine/xylazine anesthesia with a 5-mm biopsy punch. In order to limit wound closure caused by skin contraction, silicone rings were sutured to the skin around the wounds. Wounds were then topically treated with thirty microliter of AMG (diluted in DMSO, 1:3), obtained from the skin biopsy following the Rigenera^®^ protocol (*32*). A second group (n=5) was topically treated with AMG supplemented with a specific MEK inhibitor - trametinib (0.2 mg, Cat No. HY-10999, MCE). Trametinib was dissolved in dimethylsulfoxide (DMSO) to a concentration of 20 mg/ml and 10 *μ*L of the mixture (corresponding to 0.2 mg) was added topically together with the AMG (*15*). A third group (n=5) was topically treated with the MEK inhibitor - trametinib (0.2 mg, Cat No. HY-10999, MCE) (1:3 in PBS). The control group was treated with PBS diluted 1:3 with DMSO as vehicle. Wounds were covered with Tegaderm™ dressing (3M, Maplewood, USA) to protect the wound area and to prevent the area from drying out. Every other day (day 2,4, 6 and 8), digital images were taken using a Canon EOS 5D Mark II camera, treatments were re-applied topically to the wound site and dressings were renewed. Changes in % of wound closure was calculated by comparing the healed wound area of specific days with the original wound area over the time, as shown in the formula reported below (*39*) (Eq.3).

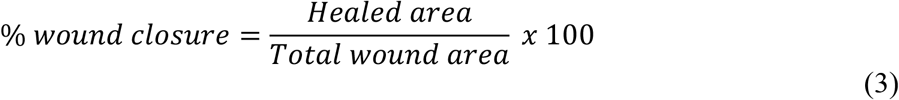

The analyses were performed using ImageJ software (NIH, Baltimore, Maryland). Animals were sacrificed at day 8 after wounding and wound skin fractions were collected to be further processed for paraffin embedding or gene expression analyses. Immunohistochemistry and immunofluorescence staining were performed on 5 *μ*m microtome sections. Animal procedures were performed in accordance with the Ethics Committee at KU Leuven (Belgium) under the ethical approval codes P056/2017 and P036/2018.

#### In vivo immunohistochemistry and –fluorescence staining

Paraffin-embedded sections were deparaffinized and rehydrated, followed by an antigen retrieval method (slides were incubated with trypsin 1:80 in 0.01% CaCl_2_ at 37 °C for 7 minutes. Samples were washed with Tris-buffered saline (TBS), incubated for 20 minutes with MeOH-H_2_O_2_ and washed again. After a permeabilization step using 0.5% triton and a wash with TBS, the samples were blocked for 45 minutes in 20% donkey serum and incubated with the primary antibody overnight at 4°C. Samples were washed with Tris-NaCl-Tween (TNT) buffer, incubated with the secondary antibody and washed again. CD31 (1:500, BD557355, Pharmingen), actin α-Smooth Muscle - Cy3™ (1:200, C6198, Sigma Aldrich), p-ERK (1:50, sc7383, Santa Cruz) and Ki67 (1:50, MA5-14520, Thermo Fisher Scientific) signals were amplified using TSA Fluorescein System (NEL701001KT, Perkin Elmer, Waltham, MA, USA), followed by secondary antibody incubation (1:300 donkey anti mouse-biotinylated IgG-B sc-2098 Santa Cruz, donkey anti rabbit-biotinylated IgG-B sc-2089 Santa Cruz) and DAB detection method. Samples were mounted using ProLong™ Gold antifade with DAPI (P36935, Invitrogen™) or nuclei were stained using Harris counterstaining and mounted using DPX. Pictures were taken with a Zeiss AxioImager Z1 Microscope and AxioVision SE64 software. Images were merged and/or quantified using ImageJ software (NIH).

#### Hematoxylin and eosin staining

Paraffin sections were heated at 57 °C for 60 minutes, deparaffinized and rehydrated. Samples were then drowned in distilled water for 5 minutes, Harris haematoxylin for 4 minutes and washed in running tap water for 2 minutes. The sections were subsequently then soaked for 1 minute each in acid alcohol, running water, blueing reagent, running water, eosin, 95% ethanol, 100% ethanol and histoclear. The slides were then mounted with DPX and left on a slide heater overnight. Pictures from paraffin sections were taken with a Leica upright microscope and the LAS. % of re-epithelialization and granulation tissue formation (expressed in arbitrary units AU) were determined using ImageJ software (NIH).

#### Sirius red staining

Sirius Red solution was prepared by mixing 0.2 g of Direct Red 80 (Sigma-Aldrich) with saturated aqueous solution of picric acid (prepared mixing 8 g of picric acid in 200 mL of distilled water). Paraffin sections were deparaffinized and rehydrated, followed by tap water wash for 10 minutes, distilled water for 5 minutes and Sirius Red solution for 90 minutes. Slides were then washed with HCl 0.01 N for 2 minutes and dehydrated with ethanol 70% for 45 seconds and twice in ethanol 100% for 5 minutes. The samples were cleared in xylol for 5 minutes (twice), mounted with DPX and left on a slide heater overnight. Pictures were taken with a Axiovert 200M motorized inverted microscope (Zeiss). Collagen quantification was determined using ImageJ software (NIH).

#### Statistical analysis

GraphPad Prism 6 was used to graph the data and GraphPad Prism software (San Diego, CA, USA) was used to analyze all the experiments of this study. Significant differences were determined by one-way or two-way analysis of variance (ANOVA) statistical tests and unpaired t test. Data are presented as mean ± standard error of mean (SEM) and as a Fold change.

## SUPPLEMENTARY FIGURES

**Fig. S1.**
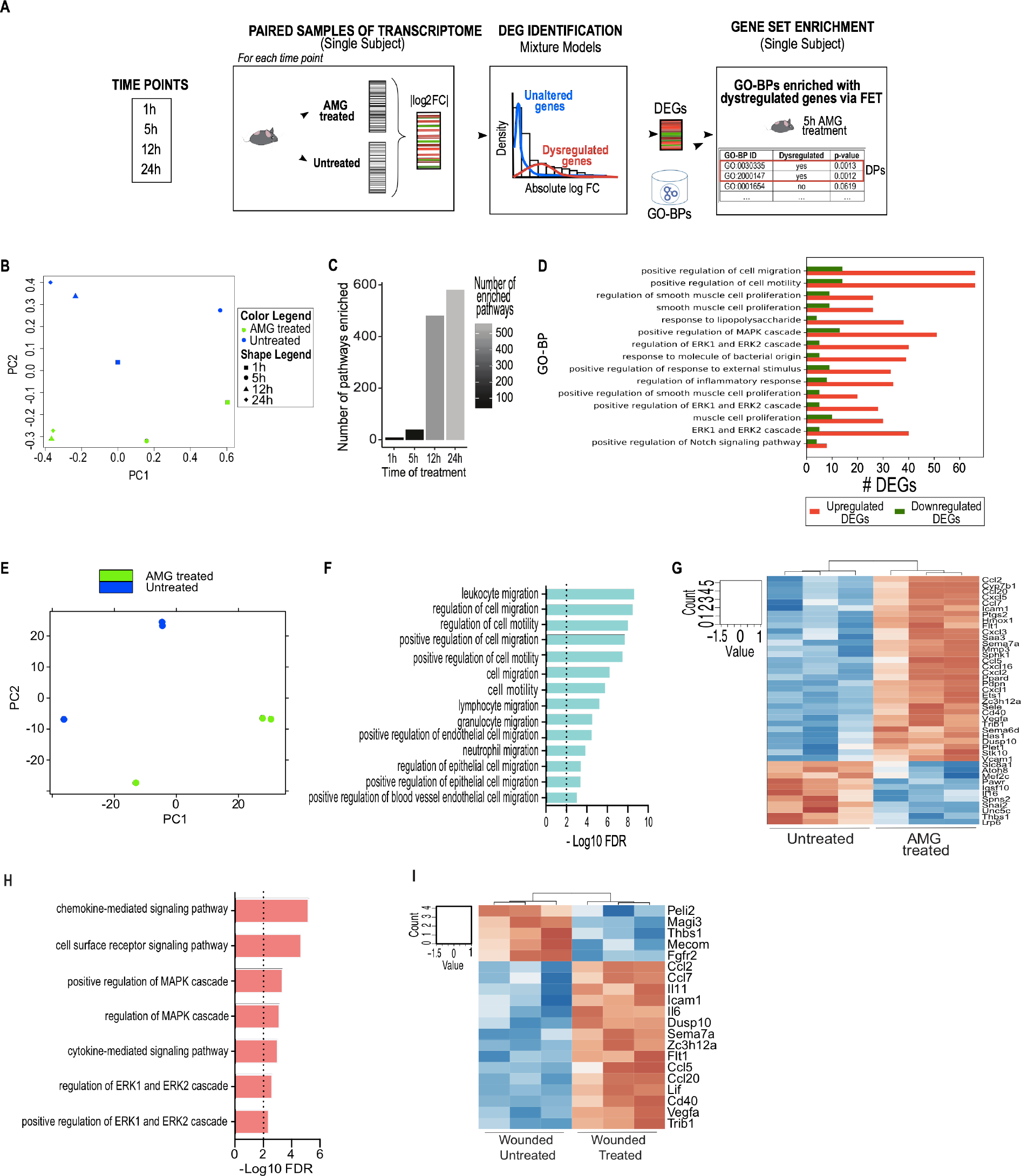
Cell migration and MAPK cascade activation as main contributors to the AMG effect. **(A)** Schematic overview of the single subject analysis performed via N-of-1 MixEnrich pathways on untreated vs AMG-treated murine primary fibroblasts. **(B)** Principal component analysis (PCA) showing transcriptome differences caused by different time of AMG treatment (1, 5, 12 and 24h). **(C)** Number of enriched GO terms associated with biological processes for each time point of AMG treatment. **(D)** Enriched dysregulated pathways upon 5h of AMG treatment obtained from Gene Ontology terms related to Biological Processes (GO-BP) (FDR<0.05). **(E)** Principal component analysis (PCA) showing transcriptome differences caused by the AMG treatment (5 hours) (3 biological replicates). **(F)** DEGs-associated enriched GO-BP selected at FDR < 0.05 and presented using GraphPad Prism 6. **(G)** Heat map showing up-regulated genes involved in process related with cell migration and cell motility (GO:0030334, GO:0030335, GO:0016477, GO:0010632, GO:0010634, GO:2000145, GO:2000147 and GO:0048870). **(H)** DEGs-associated GO terms linked with positive regulation of MAPK activity and ERK1/2 cascade were assessed as targets of the AMG treatment. **(I)** Heat map showing up-regulated genes involved in process related with ERK1/ERK2 and MAPK (GO:0070372, GO:0070374, GO:0043410, GO:0043408).

**Fig. S2.**
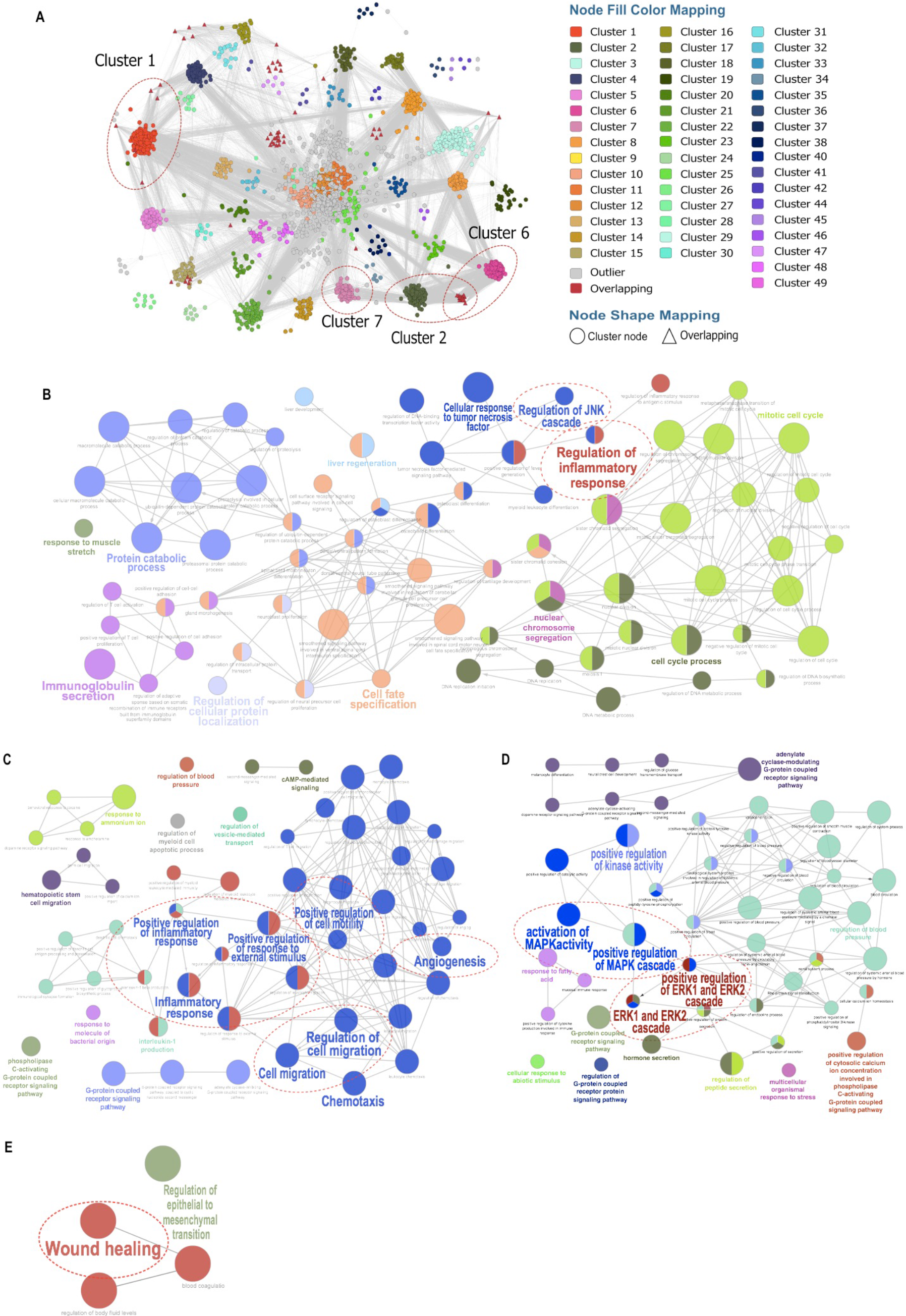
AMG treatment actively triggers important WH-associated signaling pathways. **(A)** Clusters have been obtained through ClusterONE and presented with different colors. Among the obtained 49 significant, clusters including important WH-related biological processes have been selected and presented separately. **(B)** Cluster 1: Positive regulation of the inflammatory response, including regulation of the JNK cascade, tumor necrosis factor-mediated signaling pathways as well as regulation of DNA-binding transcription factor activity have been found as candidate targets of the AMG treatment. **(C)** Cluster 2: Positive regulation of cell motility, cell migration, chemotaxis, angiogenesis and regulation of response to external stimulus have been identified as candidate target of the AMG treatment. **(D)** Cluster 6: Positive regulation of kinase activity, including positive regulation and activation of both the MAPK activity and ERK1/2 cascade has been assessed as target of the AMG treatment. **(E)** Cluster 7: Regulation of the WH process is also referred as candidate of the AMG treatment.

**Fig. S3.**
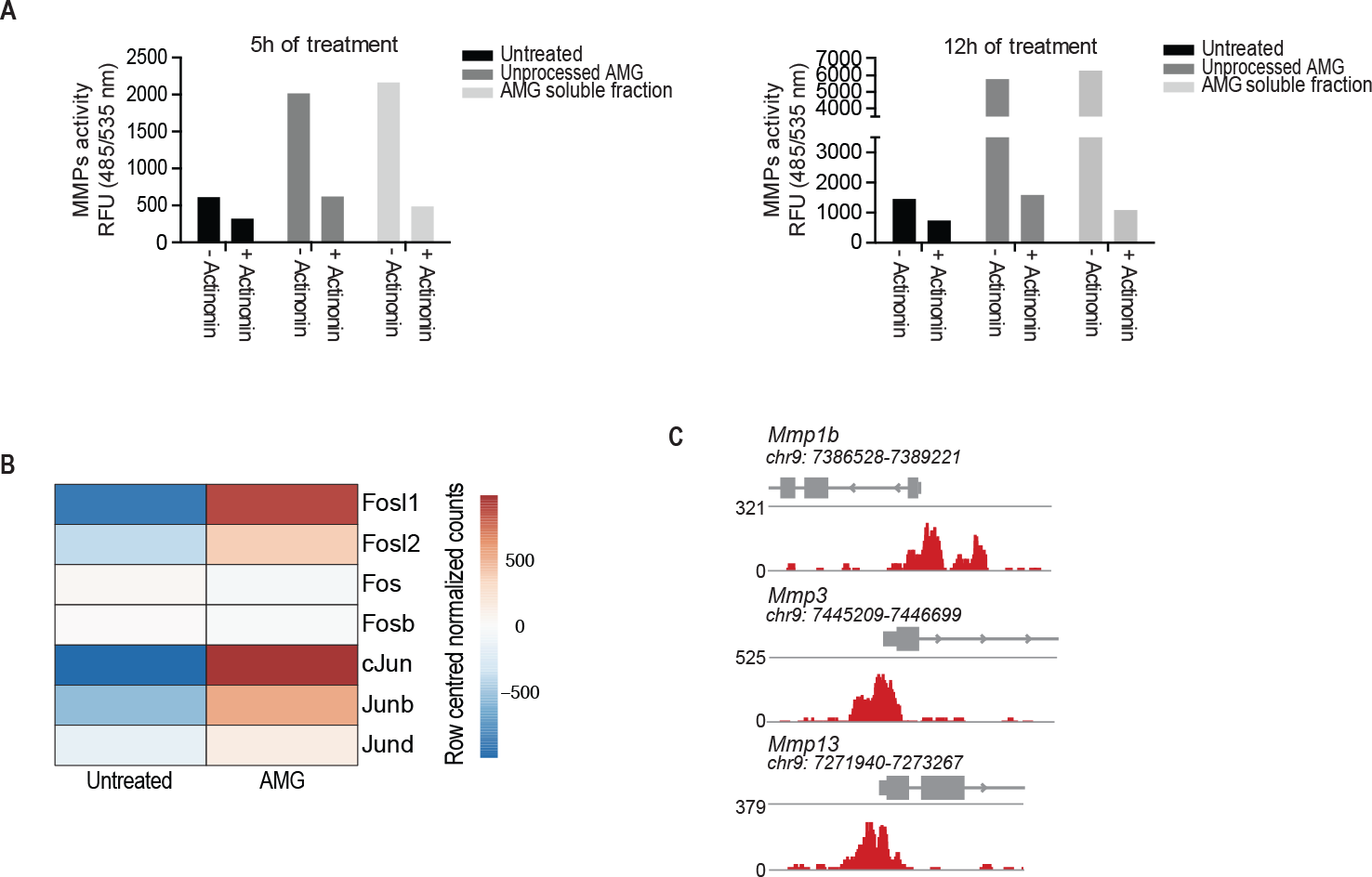
Matrix metalloproteinase enzymatic activity inhibition in an *in vitro* model of WH. **(A)** MMP extracellular activity in the biological samples exposed to the AMG treatment. Enzymatic activity of all MMP members present in cell supernatant was fluorometrically detected. Signals were evaluated 30 minutes after starting the reaction using a microplate reader with a filter set of Ex/Em = 485/535. The fluorescence signal obtained from each sample was normalized on the substrate control. Groups were incubated or not with the MMP inhibitor – Actinonin – 20µM. Groups of wounded cells that did not receive any treatment were used as a control. Data are presented as RFU (Relative Fluorescence Units) as result of technical replicates. **(B)** *In vivo* expression of AP-1 family members upon AMG treatment. Data are presented as the average of three biological replicates. (C) Levels of FRA1 binding around MMPs promoter regions.

**Fig. S4.**
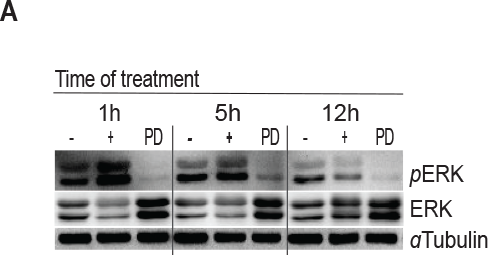
ERK phosphorylation upon AMG-based treatment in an *in vitro* model of WH. **(A)** Representative western blotting showing protein level of phosphorylated and total ERK1/ERK2 obtained from wounded murine primary fibroblasts exposed to AMG treatment and in which we induced the inhibition of the MAPK signaling pathway (using the MEK inhibitor PD0325901 – 1 µM) for different time periods (1, 5 and 12h).

**Fig. S5.**
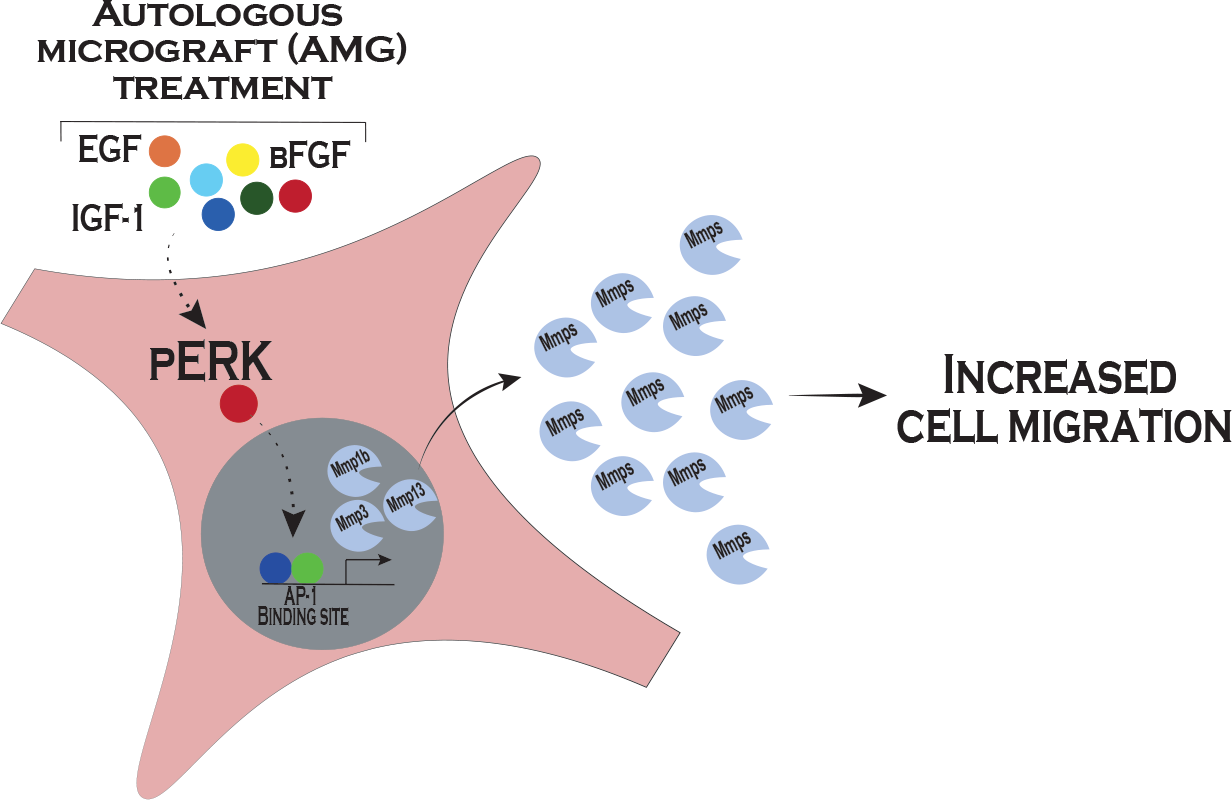
Overview of the AMG molecular mechanism. Application of AMG treatment both in *in vitro* and *in vivo* WH assays triggers to activation of both ERK signaling pathway and MMP expression and enzymatic activity. The WH-related pool of growth factors identified within the AMG may play a cardinal role in the activation of the MAPK cascade, which in turn may activate specific AP-1 family members. Activation of the transcription factor may induce MMP transcription and translation. Altogether, these events lead to an increase in cell migration rate, accelerating the whole WH process.

**Table S1.**
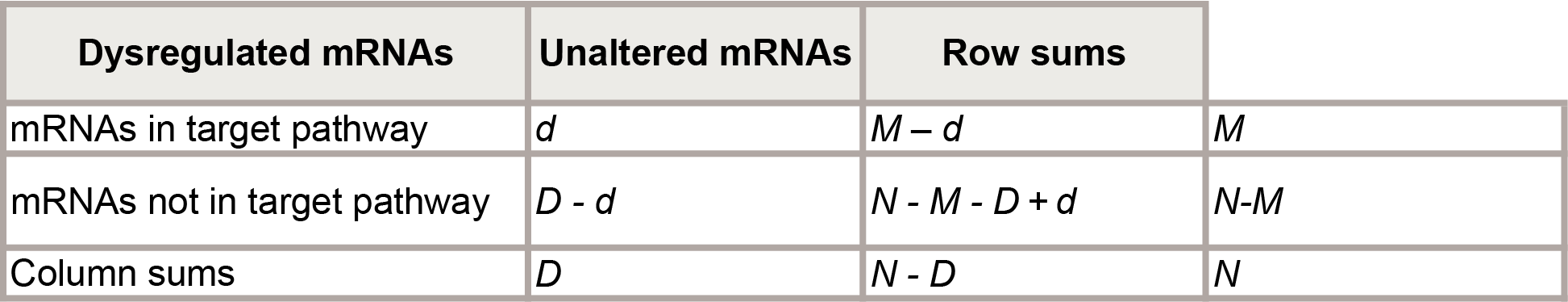
N-of-1 MixEnrich Pathways analysis upon AMG treatment.

**Table S2.**
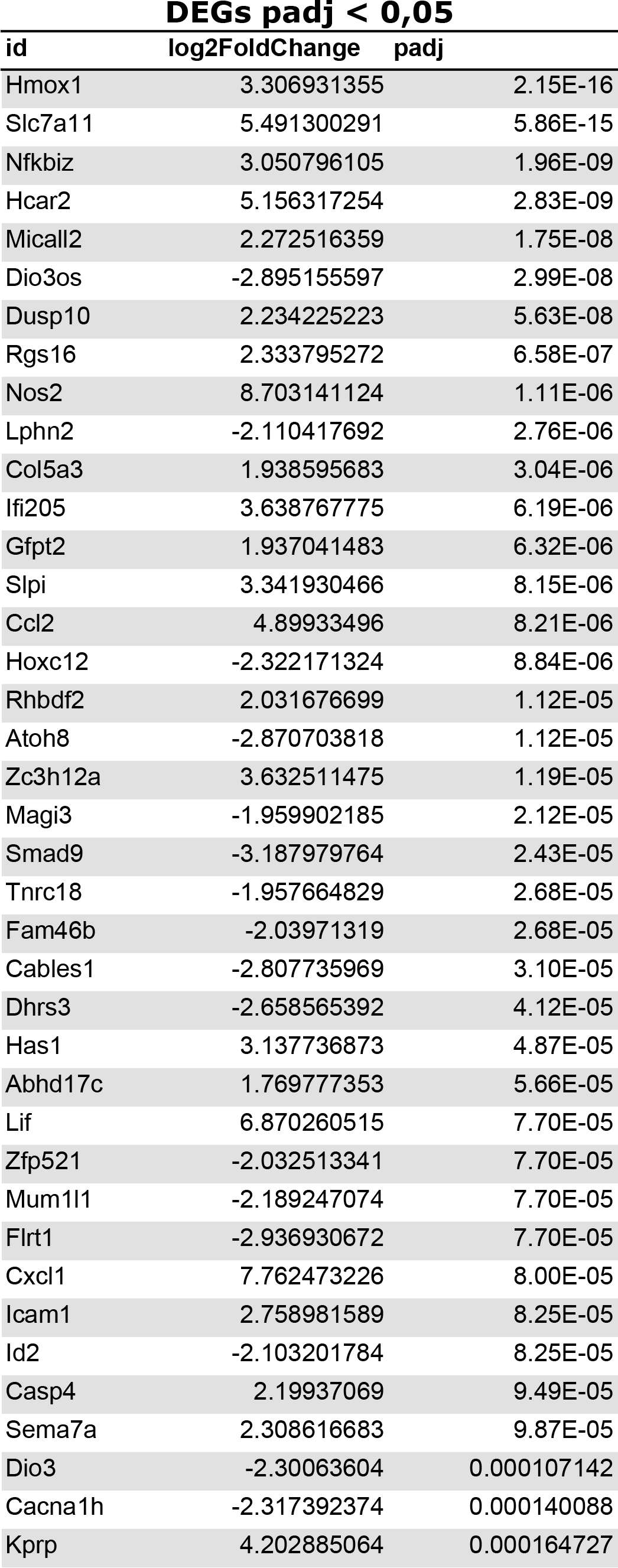
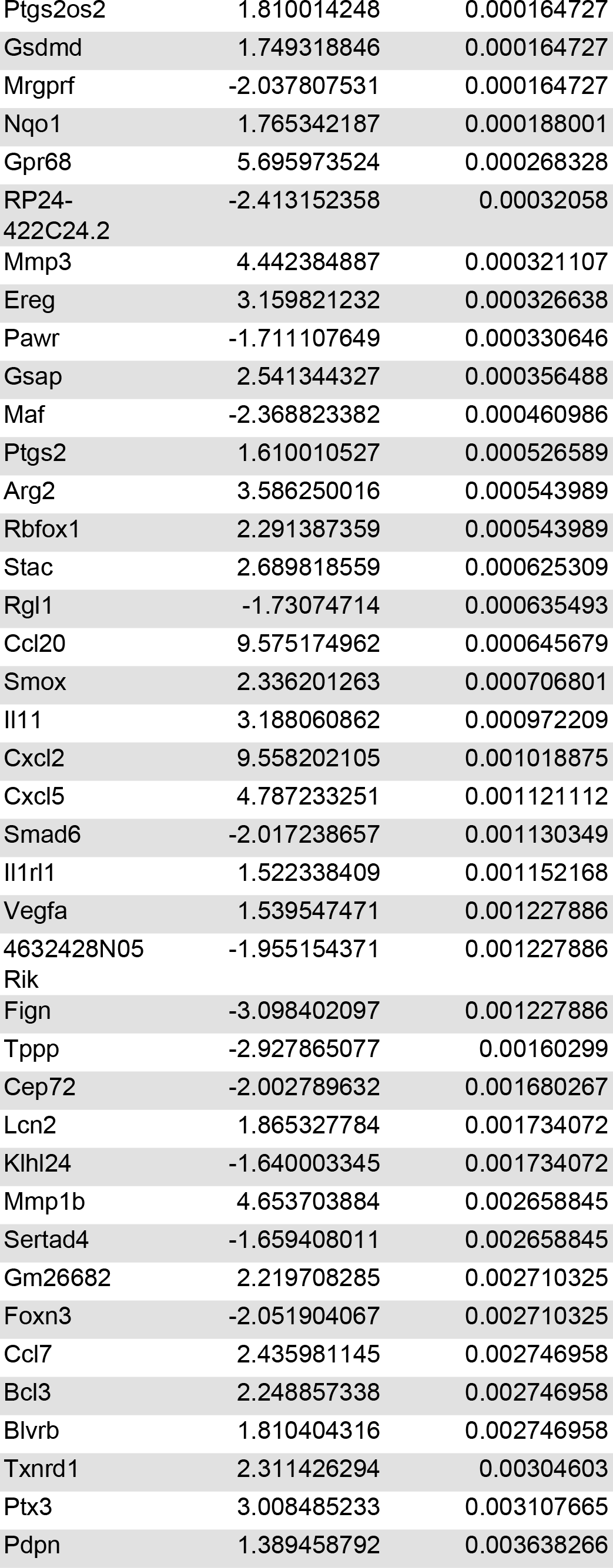
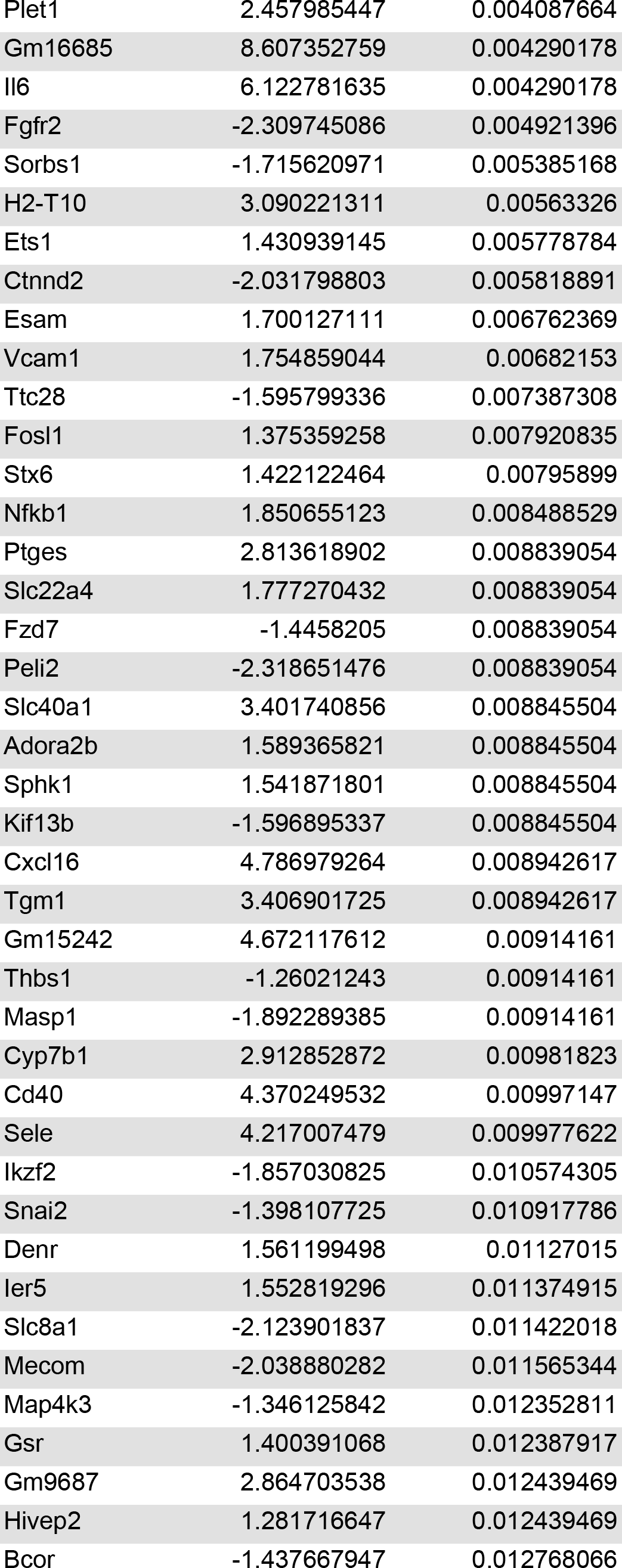
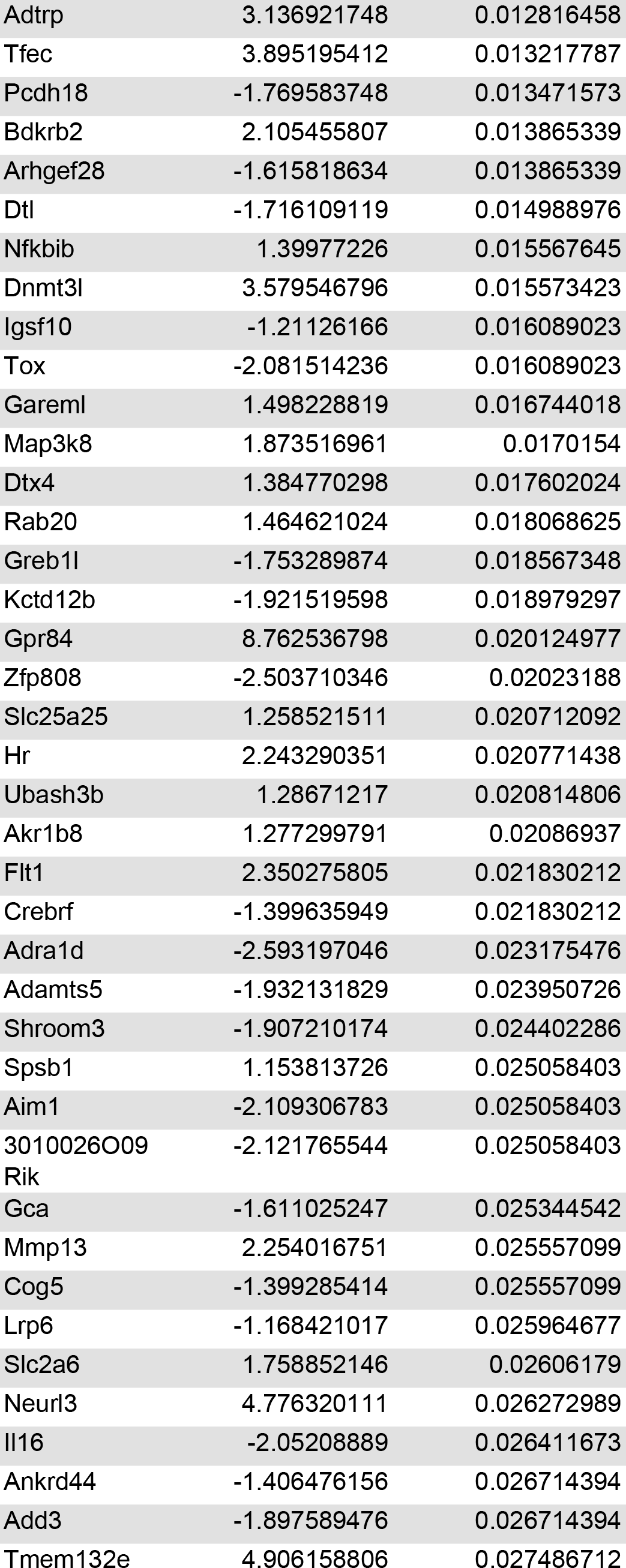
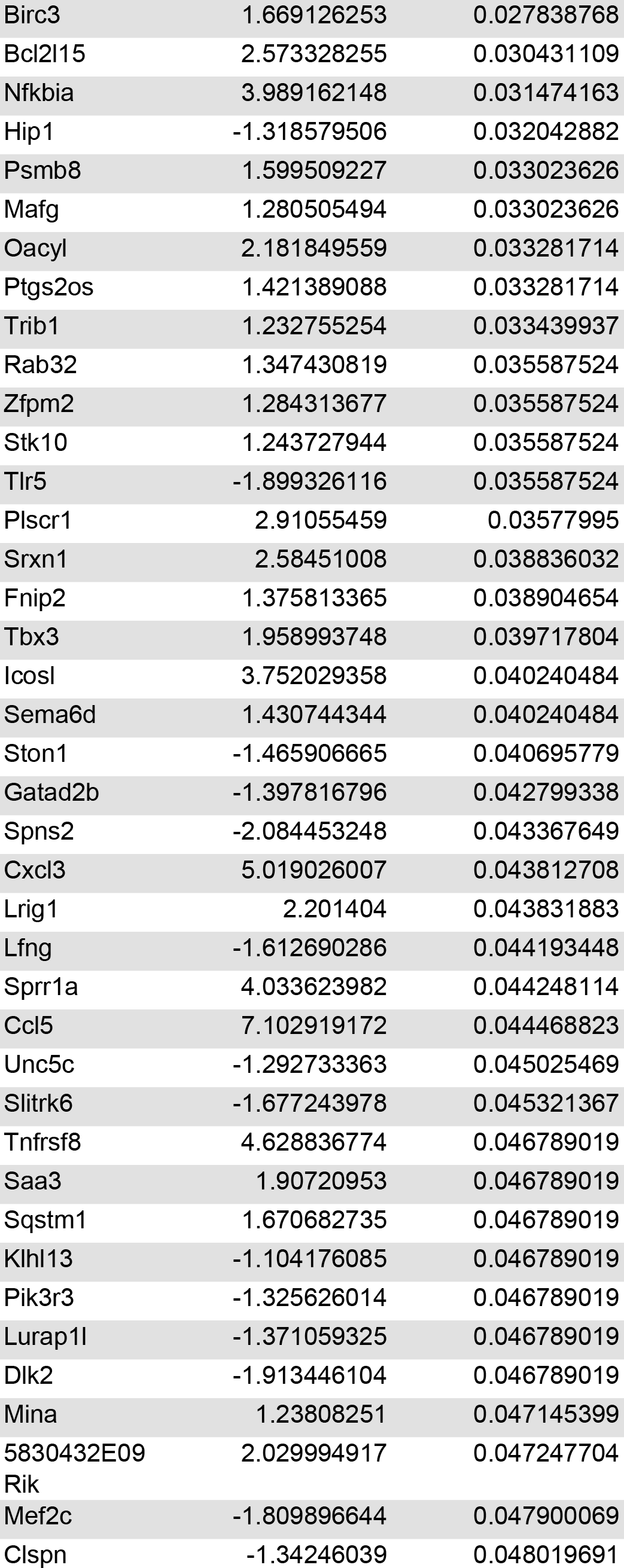
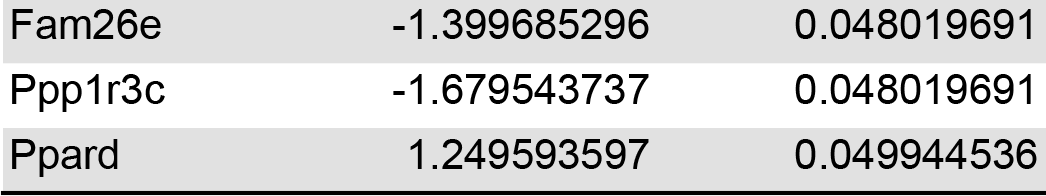
DEGs associated to AMG treatment exposure and gene ontology enrichment analysis.

**Table S3.**
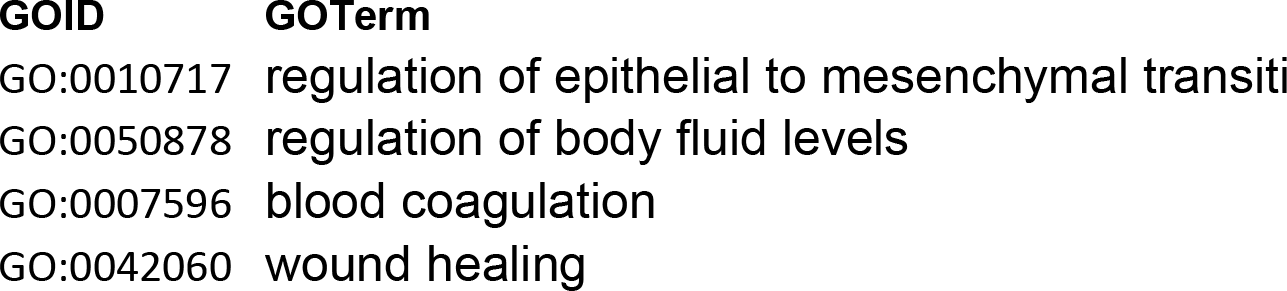
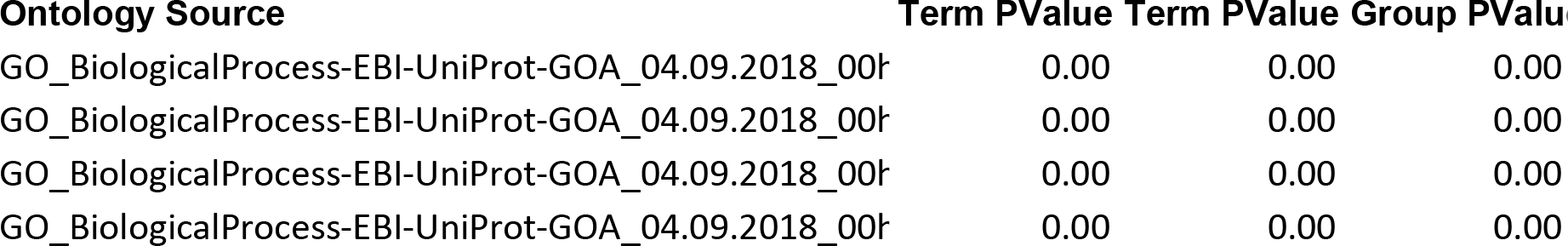
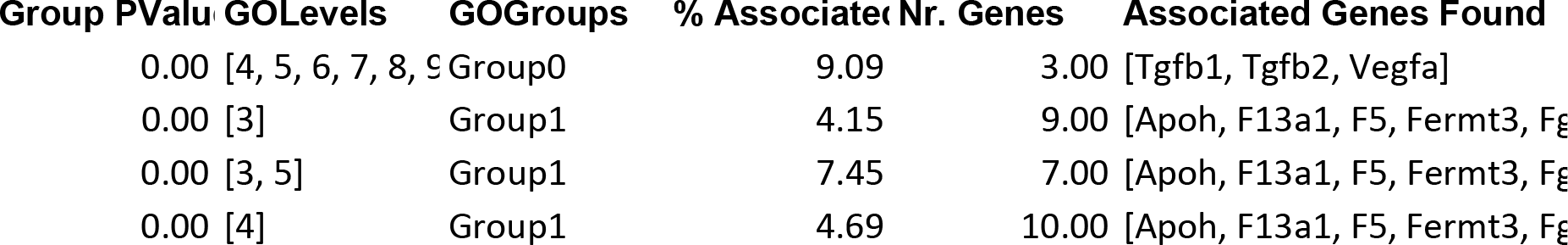
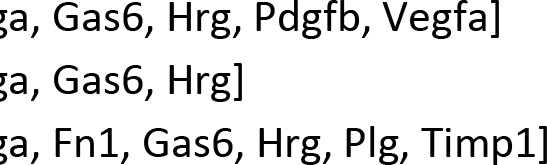
Identification of AMG targets: DEGs-related Hub nodes and cluster analysis of the AMG-mediated WH network.

**Table S4.**
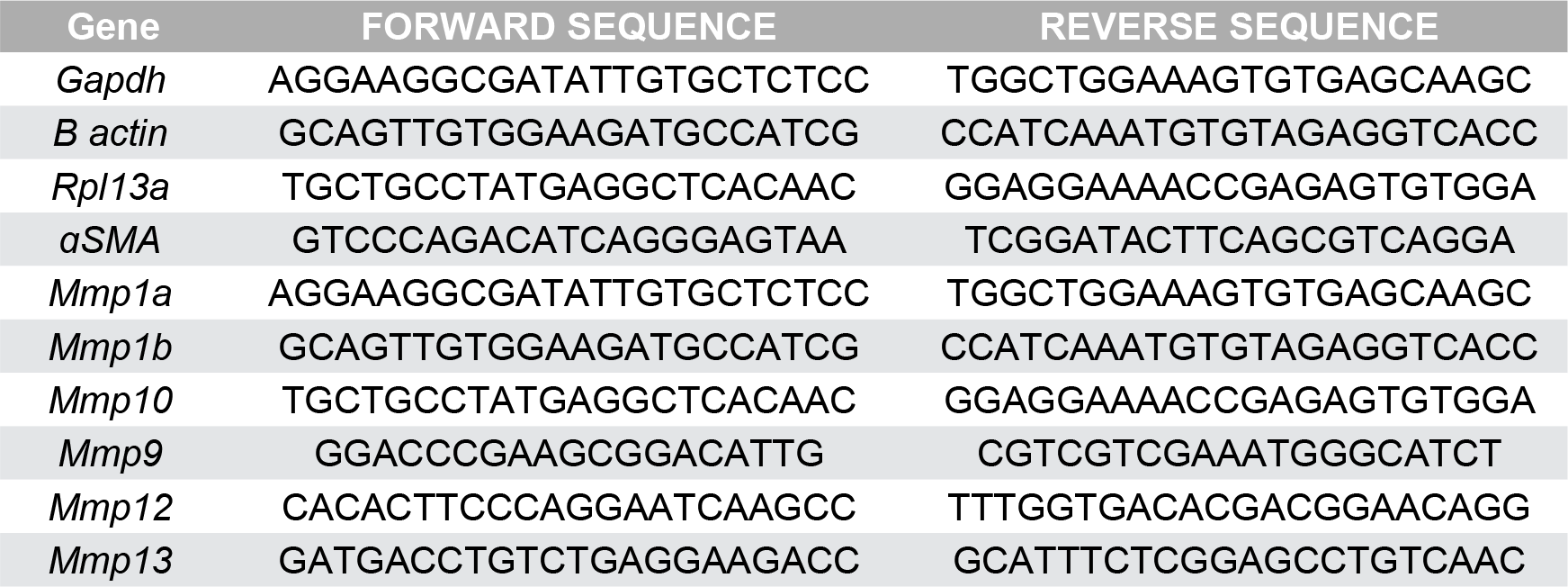
Murine Primers for qRT-PCR.

